# Unveiling the hidden biodiversity of microbial chemoreceptor genes in the rare biosphere

**DOI:** 10.1101/2025.06.24.661292

**Authors:** Claudia Sanchis-López, Saray Santamaría-Hernando, Carlos P. Cantalapiedra, Lisa Pokorny, Oscar González-Recio, Shinichi Sunagawa, Emilia López-Solanilla, Jaime Huerta Cepas

**Affiliations:** Centro de Biotecnología y Genómica de Plantas, Universidad Politécnica de Madrid (UPM) - Instituto Nacional de Investigación y Tecnología Agraria y Alimentaria (INIA-CSIC), Campus de Montegancedo-UPM, 28223 Madrid, Spain; Departamento de Biotecnología-Biología Vegetal, Escuela Técnica Superior de Ingeniería Agronómica, Alimentaria y de Biosistemas, Universidad Politécnica de Madrid (UPM), 28040 Madrid, Spain; Real Jardín Botánico (RJB-CSIC), 28014 Madrid, Spain; Departamento de Mejora Genética Animal. Instituto Nacional de Investigación y Tecnología Agraria y Alimentaria (INIA-CSIC), Ctra La Coruña km 7.5, 28040 Madrid, Spain; Department of Biology, Institute of Microbiology and Swiss Institute of Bioinformatics, ETH Zürich, 8093 Zürich, Switzerland

## Abstract

Chemoreception plays a central role in microbial adaptability, influencing both community structure and interactions with the environment. However, many chemosensitive microorganisms occur at low abundances in natural ecosystems, which has limited their detection and study using conventional metagenomic sequencing. Here, we employed a custom-designed target capture sequencing approach—encompassing all known chemoreceptor genes from both cultured and uncultured microorganisms—to uncover and characterize the vast chemosensory potential and biodiversity within the rarest fractions of the microbiome. Compared to standard environmental sequencing methods, our approach enhanced the detection of chemoreceptor (CR) genes and their associated sensing domains by orders of magnitude across diverse environments, including the rhizosphere, phyllosphere, soil, aquatic ecosystems, the human gut, and bovine rumen. This enabled the identification of thousands of low-abundance chemosensitive microorganisms that remained undetectable using conventional sequencing approaches, including known plant pathogens and symbionts. Phylogenetic analysis of the most divergent CR genes revealed evidence for novel chemosensitive species, potentially representing new bacterial phyla and classes. Our study provides a new perspective on the chemoreception capabilities of environmental microbes and opens new avenues for discovering and characterizing novel microbial sensing mechanisms.

## INTRODUCTION

Chemoperception, i.e., the ability to detect and respond to chemical cues, plays a crucial role in microbial adaptability and survival^1–3^. It regulates diverse behaviors, including nutrient acquisition^4,5^, predator evasion^6^, and interactions with hosts^7^ and other microorganisms^8^. Central to the chemosensory response are chemoreceptor proteins (CRs), which serve as primary mediators of environmental stimuli detection. These proteins, also known as methyl-accepting chemotaxis proteins (MCPs), detect chemical gradients with remarkable precision and initiate cellular responses that lead to behaviors such as motility, biofilm formation, and metabolite secretion^9–12^. In natural habitats, CRs give bacteria important advantages that are crucial for their survival^13^. Furthermore, they allow bacteria to establish symbiotic and pathogenic associations^14,15^ and strongly impact biogeochemical processes^16,17^. CRs are also relevant for establishing host-microbe interactions^14^. For instance, plant-associated microbes possess specialized CRs that detect plant-derived chemicals and guide microbes towards the plant’s roots, where both can enter into a mutualistic relationship in which the microbes boost the plant’s nutrient absorption while the plant supplies the microbes with carbon sources^18–20^. Alternatively, CRs can guide microbes towards leaf wounds and stomata where they can initiate infections^21,22^. In host-associated microbial communities, chemosensing is actively involved in biofilm formation^23,24^, playing a relevant role in feed digestion efficiency in ruminant animals^25^ and human health^26^. Similarly, chemoperception and chemotaxis are responsible for mediating the colonization process of human pathogens such as *Vibrio cholerae*, that are guided towards the intestinal mucosa^27,28^, or *Helicobacter pylori*, which colonize the mucus of the human stomach^29^.

Current knowledge on microbial chemoperception is, however, largely limited to microorganisms isolated and cultured in the lab. To date, fundamental aspects such as the extent and variability of CRs across ecosystems remain unexplored. This is partly due to the inherent difficulties of mining and classifying CR sequences obtained from metagenomics data. At the molecular level, CR genes encode for multi-domain proteins that are highly variable and encompass a large gene family. One such domain is the C-terminal methyl-accepting signal transduction domain (MCPsignal), which is highly conserved and ubiquitous among all CR genes, as it is required for the propagation of the chemosensory cascade through interactions with various kinases^30,31^. In addition, canonical CRs harbor one or more N-terminal ligand-binding domains (LBDs). These domains are extremely variable and control chemical sensing, which is why they are used to subclassify CRs according to their putative function^31,32^. Another sequence domain present in CR genes is HAMP, which plays a role in signal propagation from the LBD to the cytoplasmic signal domain^33^. Although the molecular function of HAMP may be conserved^34^, it has a low degree of sequence conservation^35^ and is not present in all CRs. Furthermore, the highly variable domain arrangement within CR sequences has led to a classification system based on CR domain topology^33,36^. Thus, the characterization and functional classification of CRs typically requires both high-quality, full-length gene assemblies and in-depth comparative analyses, which is a major bottleneck at the metagenomics scale.

Additionally, many microorganisms possessing chemosensory systems occur at low abundance in natural environments^37–39^, which further complicates their detection and genomic characterization using standard metagenomic methods. Despite achieving deep sequencing coverage, shotgun metagenomics remains heavily biased towards dominant community members, thus overlooking rare chemosensitive species. Although the biodiversity and ecological roles of these low-abundance organisms remain largely unexplored, they likely encompass rare microbes that might act as keystone species, significantly influencing ecosystem stability, resilience, and functionality^40^.

In this study, we address these challenges by developing a novel long-read target capture sequencing approach to comprehensively characterize chemoreceptor (CR) genes in environmental samples. Capture and enrichment methods are well-established techniques for selectively isolating specific genomic regions from complex DNA samples^41,42^. Our method employs an extensive, customly designed set of oligonucleotide probes that hybridize specifically to the conserved MCPsignal domain, providing a targeted sequencing approach for CR genes with nearly full phylogenetic coverage. Compared to metagenomic shotgun sequencing, this targeted approach offers substantial improvements in cost-effectiveness, sequencing depth, and ease of downstream data analysis, enabling the discovery of new chemoreceptors from low-abundance uncultured microorganisms.

## RESULTS

### A capture enrichment panel for chemoreceptor genes with broad phylogenetic coverage

Chemoreceptor (CR) genes constitute one of the largest prokaryotic gene families known to date, encompassing thousands of variants with highly diverse domain architectures. This extensive variability has so far hindered the efficient exploration of chemosensitive microorganisms in environmental samples, as sequence variability of CRs genes cannot be targeted using primer-driven PCR amplification methods. To address this limitation, we developed a custom hybridization capture panel that encompasses the full known sequence variability of MCPsignal domains, enabling targeted enrichment of CR genes from metagenomic samples.

We first established a comprehensive reference catalog of all known CR genes from both cultured and uncultured microorganisms across diverse environments. To do so, we analyzed the recent multi-habitat metagenomic dataset GMGC v1^43^, which includes 84,029 prokaryotic reference genomes and over 2.3 billion environmental genes assembled from 13,174 metagenomes spanning 14 distinct biomes. Following the approach described in Sanchis-López *et al.*^44^, we identified chemoreceptor proteins by screening for the highly conserved MCPsignal domain (Pfam: PF00015)^45^. This produced 149,326 MCPsignal-containing sequences, including 91,776 full-length CR genes, yielding an average of 7.9 CRs per Gbp of sequencing (Supp. Figure 1).

From an ecological perspective, soil and built-environment environments contributed the largest number of unique CRs to the catalog (Fig. 1a), despite representing relatively few samples in the GMGC dataset (Fig. 1b). Indeed, the rate of unique CR discovery per unit of sequencing depth (Fig. 1c and Supp. Fig. 2) was markedly lower in less biodiverse ecosystems such as the human gut microbiome (1.47 CRs/Gbp) compared to soil (18.05 CRs/Gbp) and built-environment (28.41 CRs/Gbp). Additionally, the distribution of LBD types varied across habitats (Fig. 1d), with the “unknown” LBD category being prevalent in all biomes (averaging 33%). Soil habitats showed the highest diversity of LBD types (289 different domains), followed by the human gut (238) and marine environments (211).

**Figure 1.**
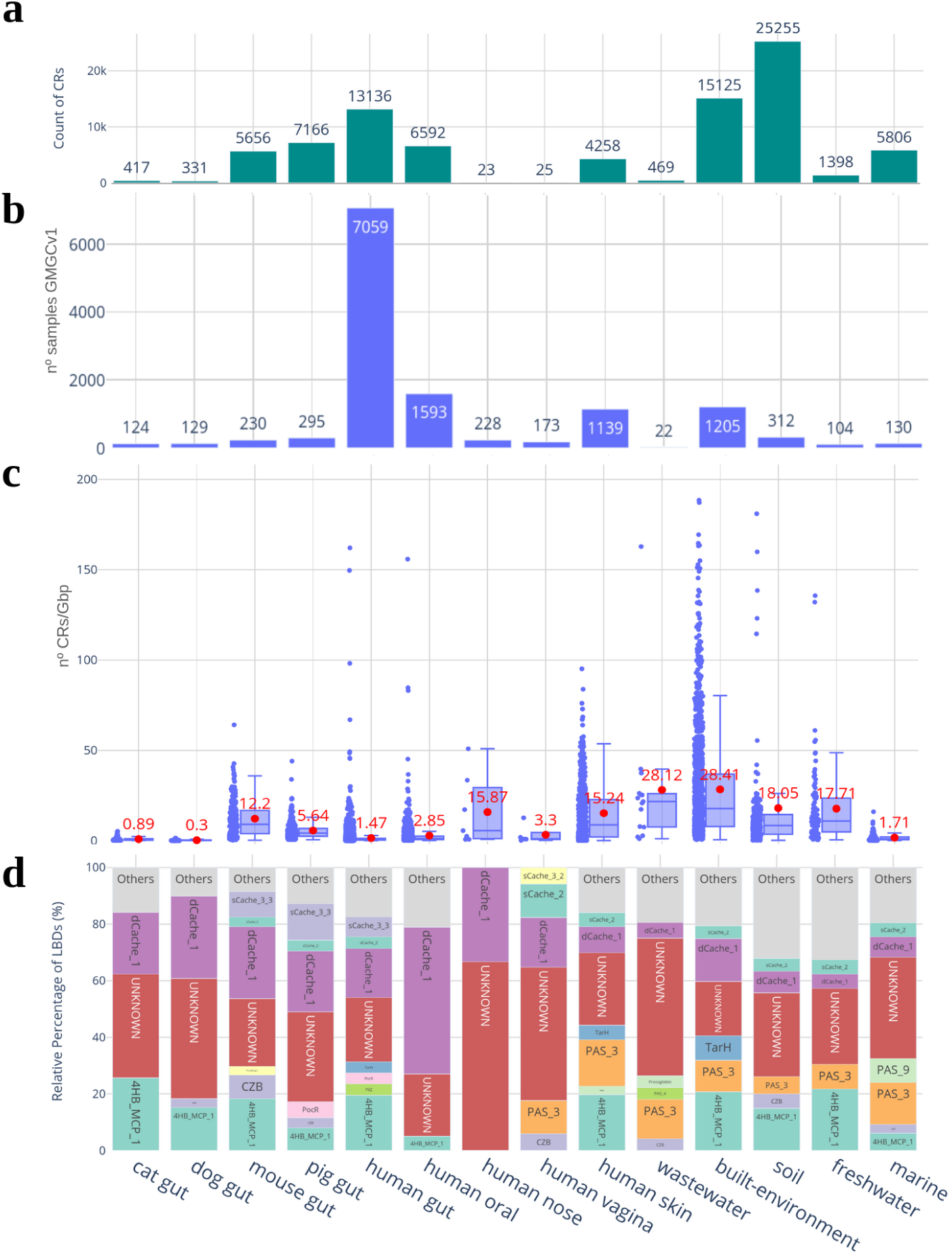
Global distribution of CRs and LBDs obtained from GMGC v1 catalog. a) Total count of CR genes retrieved from each habitat; b) Total number of samples available for each habitat; c) Normalized distribution of unique CRs per Gbp sequenced in the samples of origin per habitat, with the mean value shown in red; outliers are not shown (> 200 CRs/Gbp); d) Number of LBD sequences found per LBD-type and habitat. Domains less abundant (< 3%) are grouped as “Others”.

The resulting global CR catalog was then used to build a custom capture panel covering the entire sequence variability of CR genes observed in cultured and uncultured microorganisms across habitats. Specifically, we extracted the MCPsignal domain sequence of all 149,326 observed CRs and generated 80-nucleotide (nt) probes from them, using a sliding window of 20 nt. This approach yielded a total of 3,429,805 probes, which were then clustered at 80% identity to minimize redundancy, resulting in 1,323,163 cluster-representative probes. The final probe set was synthesized as a single Roche NimbleGen EZ Developer DNA probe set.

### Target capture sequencing boosts CR gene identification in environmental samples

To evaluate the efficiency of our CR capture panel across various environments, we performed two comparative analyses between target capture sequencing and conventional shotgun metagenomics.

Firstly, we tested our method on a diverse pool (Pool-1) of 14 samples covering the rhizosphere, phyllosphere, bulk soil, lake water, human gut, and bovine rumen environment. Following standard metagenomic sequencing procedures, we extracted DNA from each sample and sequenced it using Illumina PE150 technology for a total output of approximately 8 Gbp per sample. Concurrently, DNA from the same samples was fragmented to an average length of 1 Kb to produce a single pool of genomic libraries (Pool-1, Supp. Table 1) and processed with our custom capture panel prior to sequencing (Fig. 2). Products derived from the capture experiments were sequenced using Oxford Nanopore Technologies (ONT). To accommodate multiplexed long-read target capture sequencing, we developed a custom protocol, inspired by Bethune *et al.*^46^, which integrates Illumina-based indexing with ONT genomic library preparation and sequencing (details in the Methods section). This was necessary as ONT multiplexing kits cannot yet be adapted to standard capture protocols^47^. CR genes from both sequencing methods were assembled and deduplicated to account for unique CR counts. To mitigate potential biases from the higher error rates in ONT reads, we clustered capture-based CR genes at 90% nucleotide identity. Results highlight large differences in the number of unique CRs identified by capture-based experiments compared to conventional metagenomics across all environments, both in absolute number per sample (Fig. 3a) and normalized CRs counts per sequencing unit (Fig. 3b). Overall, we observed an increase in the number of unique CR genes detected by capture-based results compared to regular shotgun data, ranging from 2.6-fold increase in lake water samples, to 282-fold in phyllosphere samples.

**Figure 2.**
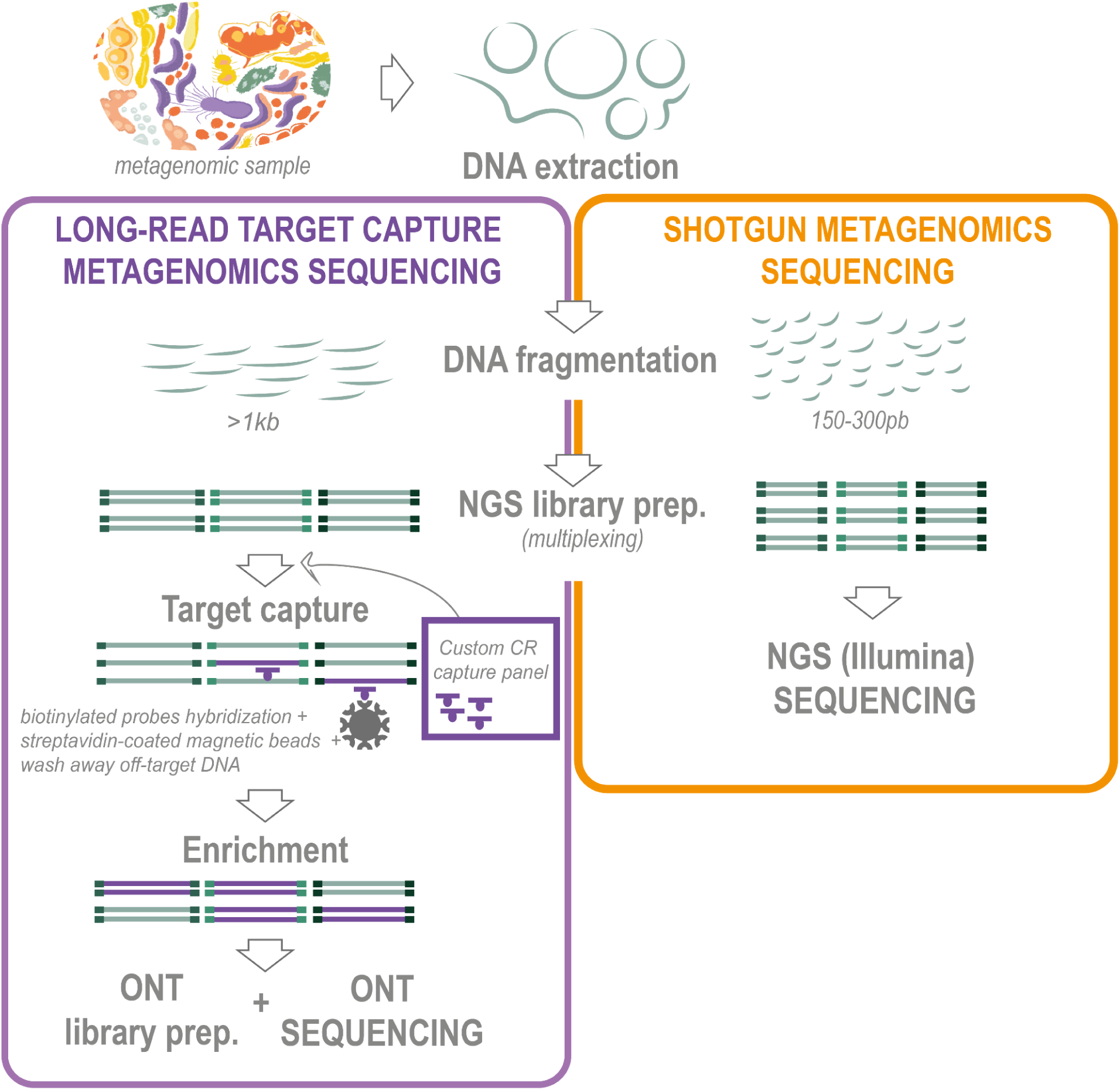
Schematic representation of the target capture sequencing approach vs standard metagenomics sequencing. Long-read target capture metagenomic sequencing (left, purple) involves long DNA fragments (∼1kb - 1.5kb), library preparation, and hybridization against the global MCPsignal-based capture panel developed in this study. Captured DNA is then amplified, and sequenced using Oxford Nanopore Technologies. Shotgun metagenomics (right, orange) involves short-read DNA fragmentation (∼300 bp), Illumina-compatible library preparation, and paired-end sequencing.

**Figure 3.**
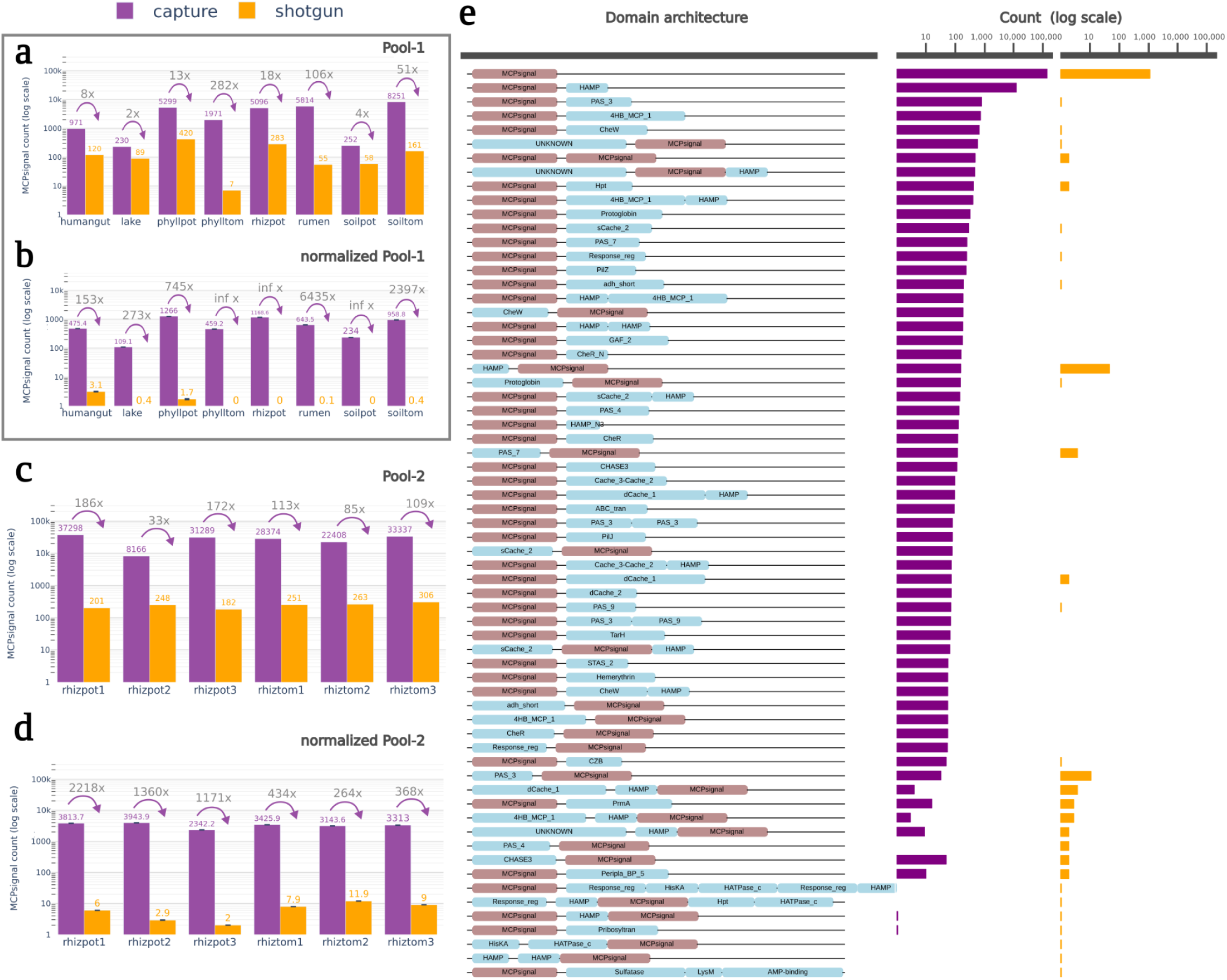
Long-read target capture metagenomic sequencing approach boosts detection of CRs across diverse samples. Total count of unique a) MCPsignal domains detected in Pool-1; b) MCPsignal domains detected in Pool-1, normalized by sequencing depth at 40 Mbp (average of 10 randomized sets for Pool-1); c) MCPsignal domains detected for each sample in Pool-2; d) MCPsignal domains detected in Pool-2 after normalization at 374 Mbp (average of 10 randomized sets for Pool-2); and e) the most frequent domain architectures detected in CR sequences obtained in Pool-2.

Secondly, we used capture-based sequencing on six ecologically homogeneous rhizosphere samples (Pool-2). This approach aimed to minimize the sequencing imbalances observed in Pool-1, likely due to the heterogeneity of samples mixed in the same pool. In this case, DNA was fragmented to 1.5 Kbp, trying to achieve longer read sequencing compared to Pool-1. Overall, target capture sequencing of Pool-2 significantly enhanced CR gene discovery in the rare biosphere by various orders of magnitude compared to shotgun metagenomics. After assembling, deduplicating and clustering CR sequences below the ONT average sequencing error, targeted sequencing on rhizosphere samples yielded a total of 160,872 unique CRs, leading to an average increase of 116-fold in the absolut number of unique CR genes identified per sample (Fig. 3c) and a 1,954-fold improvement in the normalized count of CR genes per sequencing unit (Fig. 3d). Apart from quantitative improvements, the CR genes obtained from target capture sequencing in Pool-2 were significantly longer than those assembled from regular shotgun metagenomic data (1,300 nts vs. 521 nts). This increased length translated into greater efficiency in identifying the sensing Ligand Binding Domains (LBDs) associated with CRs, which are typically located near the gene’s N-terminus and are challenging to obtain from short gene assemblies (Fig. 3e). Consequently, target capture sequencing of Pool-2 enabled us to identify 9,918 CR genes with at least one known LBD, a potential sensing domain, while bulk metagenomics sequencing identified only 63 CR gene assemblies with known LBDs.

### Unveiling the hidden biodiversity of low-abundance chemosensitive microorganisms

To explore the phylogenetic biodiversity and potential functional variability unveiled by capture-based CR genes, we mapped all CR sequences observed in Pool-2 samples (rhizosphere) against the Genome Taxonomy Database r207 (GTDB)^48^, a comprehensive resource comprising 65,703 reference bacterial and archaeal species derived from 317,542 genomes and metagenome assembled genomes (MAGs).

Using BLAST, we searched for identical matches between the MCPsignal domain sequence of CR genes, and the reference genomes or MAGs included in GTDB r207. To ensure accurate taxonomic annotations and avoid possible misassignments, we called the presence of a species only if an identical (100% nucleotide identity) and unambiguous (second-best hit <100% nucleotide identity) BLAST hit was found. Under these stringent criteria, 1,063 capture-based CR genes found perfect coincidences, putatively identifying 307 distinct GTDB species (Table 1) from 16 bacterial and 2 archaeal phyla (Fig. 4, purple bars). In contrast, conventional shotgun sequencing yielded taxonomic annotations for only 12 CR genes from six species (Fig. 4, orange bars). Of all species detections derived from capture-based data, 135 were supported by two or more CR identical matches per genome, whereas only two species met this threshold in the shotgun-based dataset (Fig. 5a). This underscores the effectiveness of our target capture sequencing approach in revealing the hidden biodiversity of low-abundance chemosensing microorganisms within highly complex environments.

**Figure 4.**
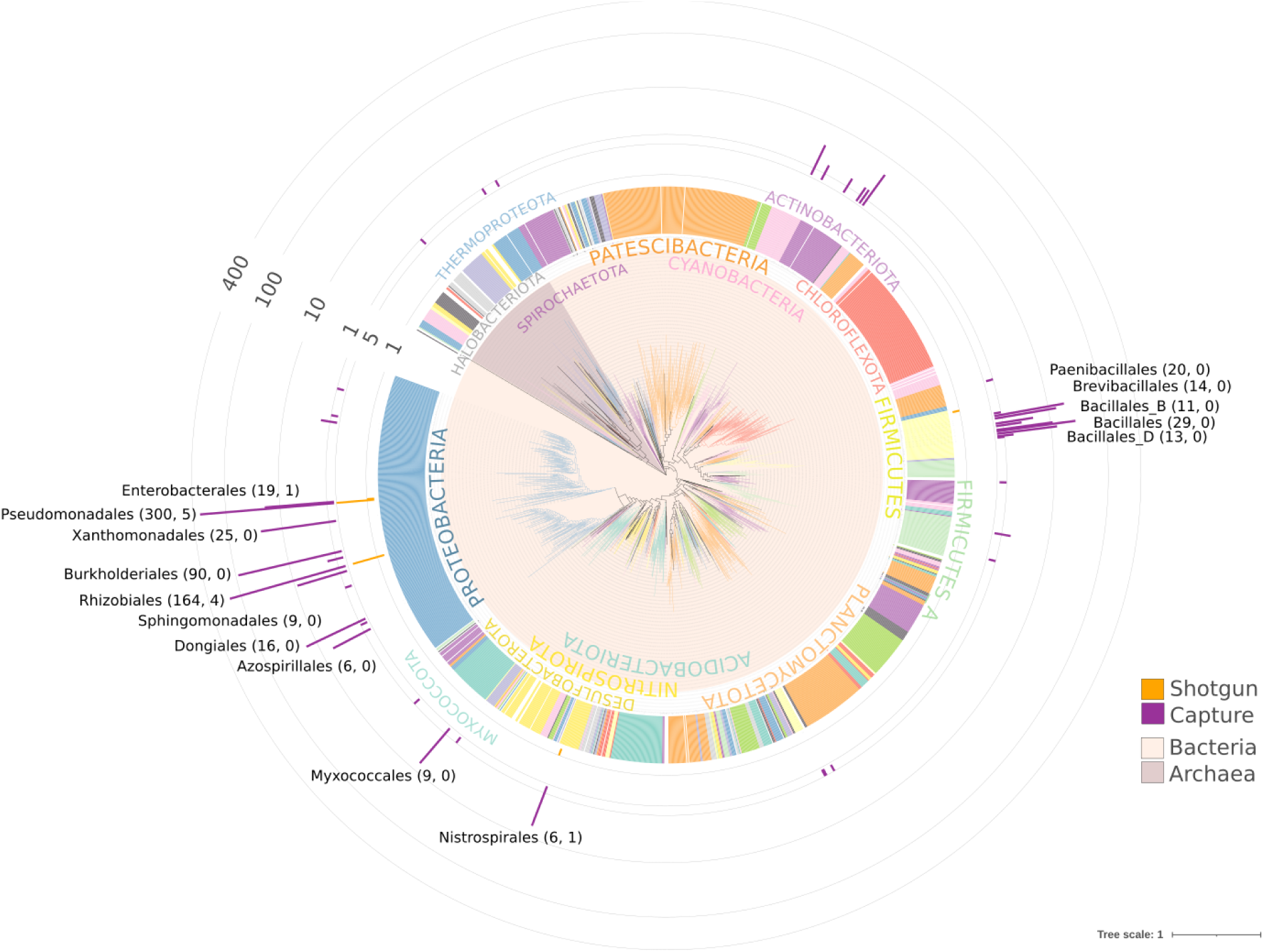
Phylogenetic distribution of taxonomic identifications achieved by identical and unambiguous CR gene mapping against GTDB genomes. The underlying tree shows GTDB r207 archaeal and bacterial phylogeny collapsed at order level. The number of CR sequences confidently assigned to a known species in GTDB within each taxonomic order based on capture-based data is shown as purple bars in log-scale, while shotgun-based CR matches are shown in orange. Taxonomic orders with more than 5 CRs are indicated, with the number of CRs indicated in parenthesis for capture-based and shotgun-based respectively.

**Figure 5.**
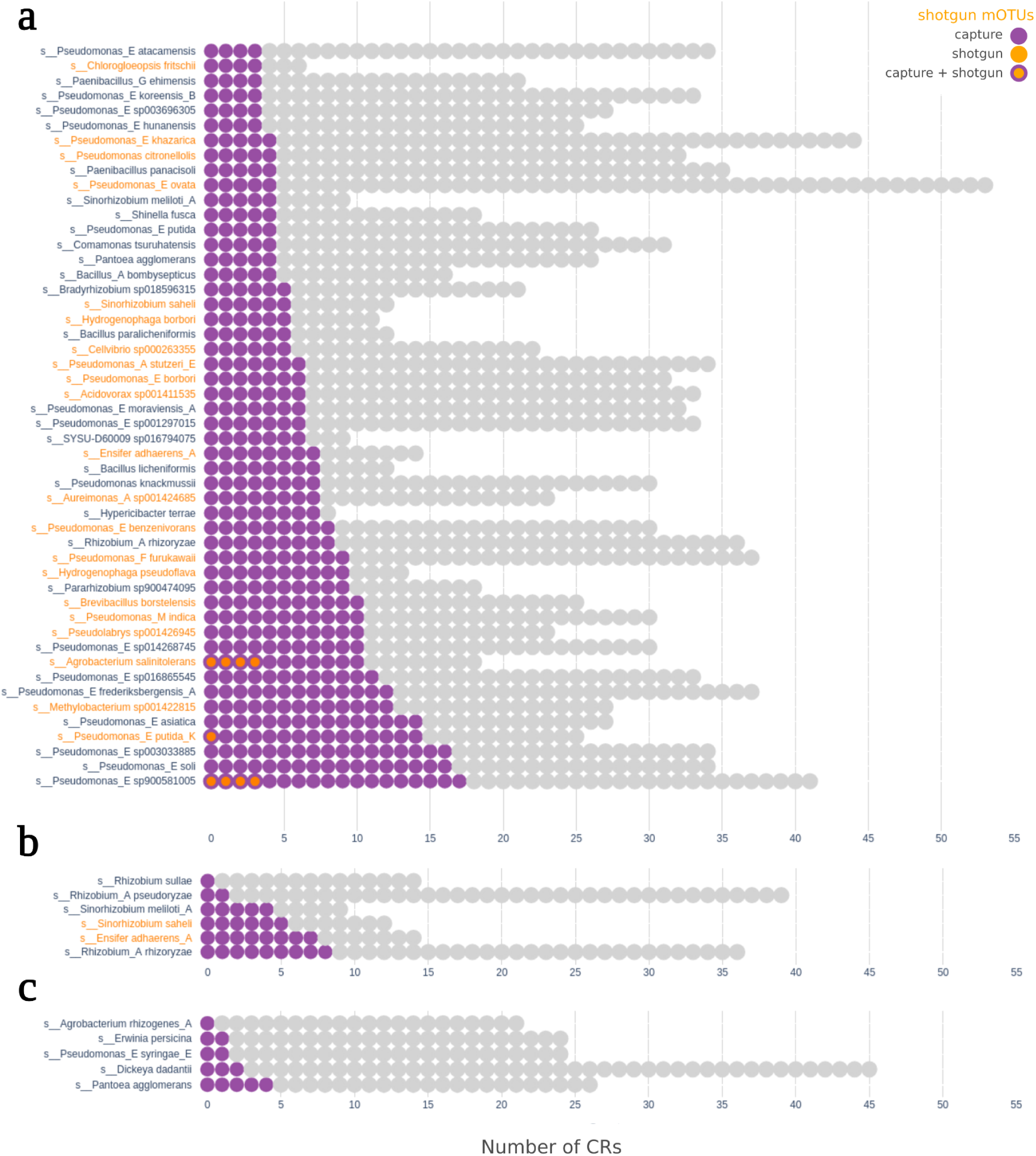
Species identification based on multiple unambiguous CR mappings. Species names in GTDB nomenclature are shown as rows with an array of circles representing their known repertoire of CR genes. Species-specific CR genes identified only by capture-based data are shown as purple circles in each array. CRs identified by both capture- and shotgun-based data are shown as purple-orange circles. Undetected CRs are shown in gray. The names of species identified independently by the mOTUS species profiler tool are indicated in orange. The panels separate species based on their classification as a) commensals, b) plant symbionts, or c) plant pathogens.

**Table 1.**
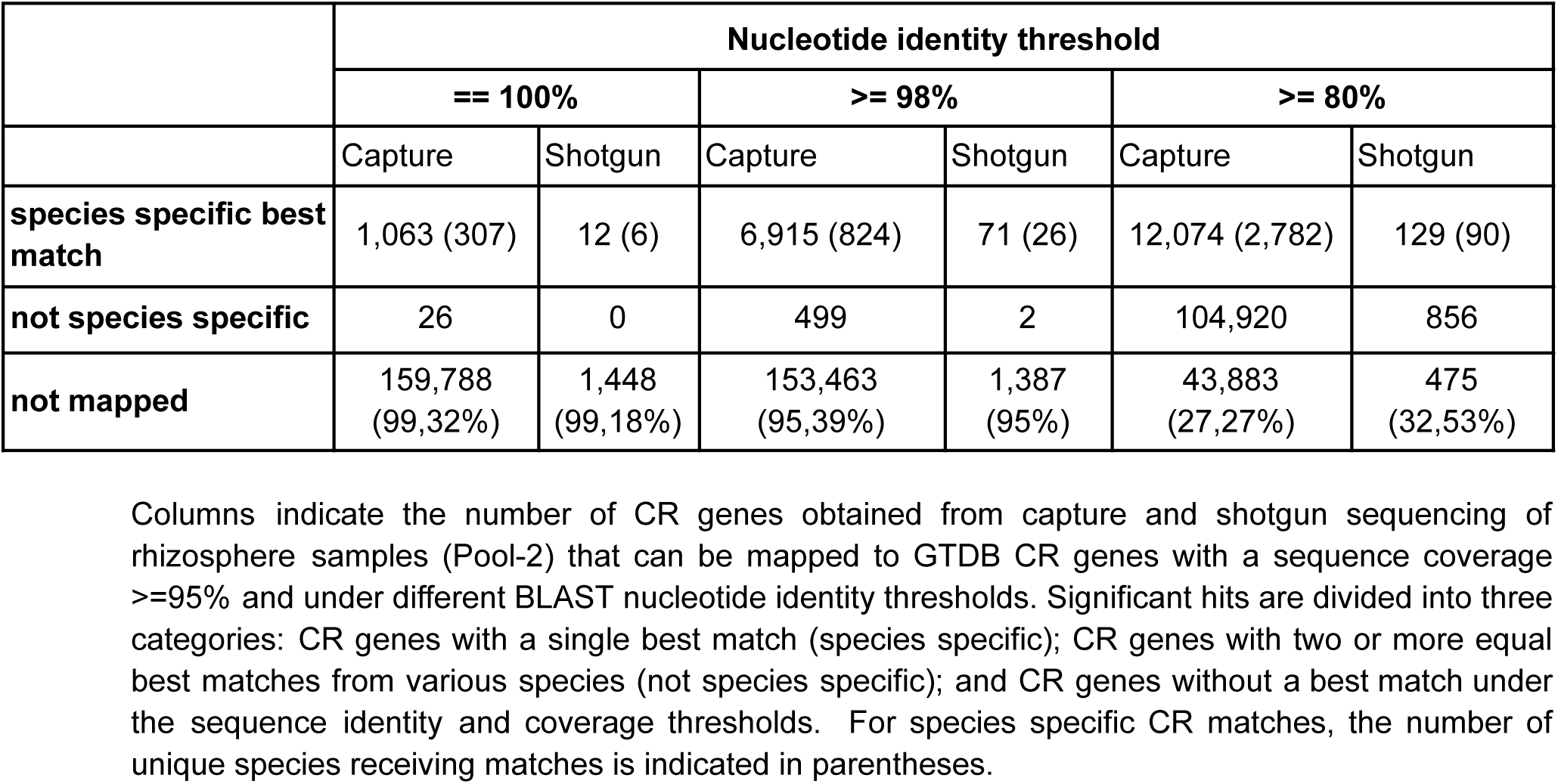
Number of capture and shotgun-based CR sequences from Pool-2 whose MCPsignal domain can be mapped to GTDB known CRs under various nucleotide identity thresholds.

|  | Nucleotide identity threshold |  |  |  |  |  |
| --- | --- | --- | --- | --- | --- | --- |
|  | == 100% |  | >= 98% |  | >= 80% |  |
|  | Capture | Shotgun | Capture | Shotgun | Capture | Shotgun |
| **species specific best match** | 1,063 (307) | 12 (6) | 6,915 (824) | 71 (26) | 12,074 (2,782) | 129 (90) |
| **not species specific** | 26 | 0 | 499 | 2 | 104,920 | 856 |
| **not mapped** | 159,788<br>(99,32%) | 1,448<br>(99,18%) | 153,463<br>(95,39%) | 1,387<br>(95%) | 43,883<br>(27,27%) | 475<br>(32,53%) |

Columns indicate the number of CR genes obtained from capture and shotgun sequencing of rhizosphere samples (Pool-2) that can be mapped to GTDB CR genes with a sequence coverage $\geq 95\%$ and under different BLAST nucleotide identity thresholds. Significant hits are divided into three categories: CR genes with a single best match (species specific); CR genes with two or more equal best matches from various species (not species specific); and CR genes without a best match under the sequence identity and coverage thresholds. For species specific CR matches, the number of unique species receiving matches is indicated in parentheses.

Notably, target capture sequencing identified 11 known plant pathogens and symbionts, whose low abundance prevented their detection using standard sequencing methods (Figs. 5b and 5c). Key examples include the pathogens *Pantoea agglomerans, Dickeya dadantii,* and *Pseudomonas syringae*, supported by the concurrent unambiguous detection of multiple CR genes in capture-based data (purple dots in Figure 5), but undetected by the CR genes from shotgun data (orange dots, in Fig. 5). Interesently, these pathogens were also undetected when using mOTUs^49^, a standard shotgun-based species profiling method based on phylogenetic marker genes (mOTUs detections indicated with orange names in Fig 5). Similarly, known plant symbionts such as *Ensifer adhaerens, Rhizobium rhizoryzae, Sinorhizobium saheli,* and *Sinorhizobium meliloti* were identified exclusively through the capture-based approach. In total, only two out the 11 latent microorganisms were detectable using bulk metagenomics data, either by CR mapping or through mOTUs taxonomic profiling.

Regarding the functional diversity of CR genes, the overall longer capture-based CR sequences unveiled a much larger repertory of putative sensing domains than those assembled from bulk metagenomic data, suggesting a vast hidden repertory of sensing domains in CRs from low abundance species. Following a previously validated procedure^44^, we extracted all putative ligand-binding domains (LBDs) present in both capture-based and shotgun-based CR genes. In total, we identified 10,533 putative LBDs from 9,918 captured-based CR genes, but only 215 from shotgun data. Besides quantitative differences, capture-based data also uncovered many novel putative sensing domain families. Out of all 10,533 LBD sequences detected, 2,242 (21.3%) had no homologs in GTDB, and 577 of these could not be matched to any previously described Pfam domain (Supp. materials), suggesting an extensive range of new sensing capabilities present in rare uncultured taxa.

### Novel chemosensitive species in the rare biosphere

A significant proportion of capture-based CR genes in Pool-2 could not be confidently mapped to known genomes or MAGs in GTDB (see Table 1). In fact, 27% of all captured-based CR genes contained a MCPsignal domain sequence below 80% of nucleotide identity to their best GTDB match. This suggests the presence of a large number of unknown chemosensitive species in the rare biosphere that could potentially be identified by their CR repertoire.

We therefore reconstructed a comprehensive phylogenetic tree containing all the MCPsignal domain sequences from CRs present in GTDB genomes and MAGs, alongside those from capture-based CR genes. As expected, most capture-based CR genes grouped closely within known species and genus clades. However, we identified 38 highly divergent and well-supported tree branches grouping 100 or more novel CR sequences monophyletically (Supp. Table 2). Seven of these capture-only clades had an estimated last common ancestor at the domain taxonomic rank, and contained CRs whose MCPsignal’s best match in GTDB had an average protein identity of less than 70%. Although the MCPsignal domain is not a robust phylogenetic marker due to its multicopy nature and tendency for horizontal gene transfer, the consistent divergence observed across entire clades of novel CRs may indicate vertical diversification from ancient and unknown lineages. These lineages are likely to represent new phyla or classes of chemosensitive microorganisms. For example, clade N330390 (see Fig. 6a) comprises 121 putative new CR sequences and exhibits an average protein identity of 54.7% with all known cultured and uncultured prokaryotic species. It is also phylogenetically placed between CRs of various phyla, including *Synergistota, Firmicutes, Proteobacteria, Patescibacteria, Nanoarchaeota* and *Goldbacteria* (see Fig. 6a). Similarly, clade N231481 (with an average protein identity of 56.9%) groups together 113 highly divergent CR sequences and is placed between CR sequences from the *Asgardarchaeota*, *Myxococcota* and *RBG-13-61-14* phyla (Fig. 6b).

**Figure 6.**
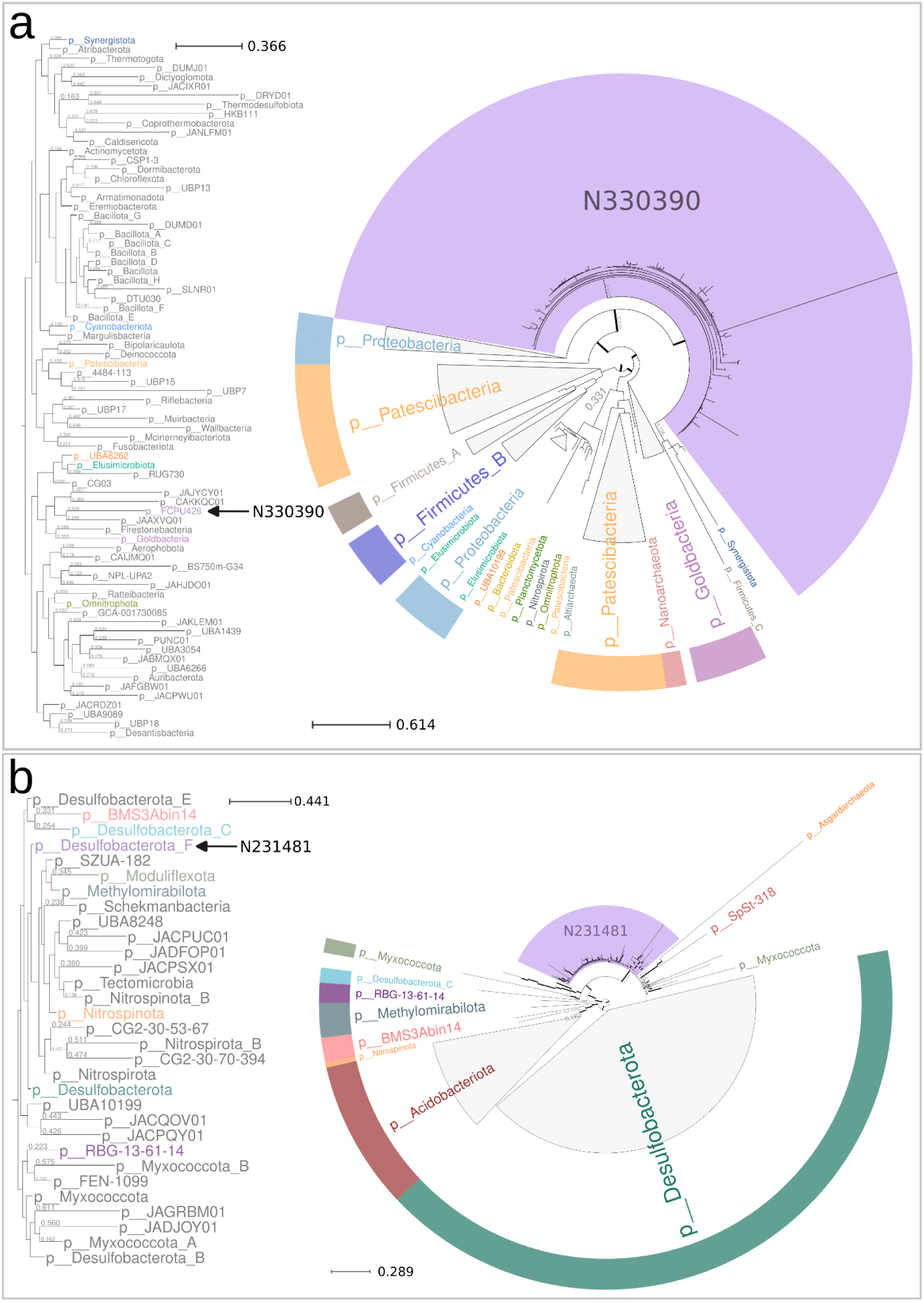
Phylogenetic placement of two highly divergent sets of CR sequences uncovered by target capture sequencing. The circular trees represent subparts of the global MCPsignal domain phylogeny reconstructed in this study and available as Supp. material. Panels show two subtrees containing the clades N330390 (purple background, panel a) and N231481 (purple background, panel b), alongside with the phyla classification of their closest branches. The rectangular trees on the left of each panel show the taxonomic classification of the best matching MAGs of each CR clade in the SPIRE database (marked with arrows), in the context of the GTDB reference species tree.

To confirm that these divergent CR clades in the MCPsignal tree might indicate highly divergent novel chemosensitive species, we performed a systematic search for such novel CRs against the Searchable Planetary-scale Microbiome Resource (SPIRE)^50^, which provides a consolidated set of more than a million metagenome-assembled genomes (MAGs), assembled from 99,146 metagenomic samples. As well as providing an updated database of observed uncultured species, SPIRE provides taxonomic annotations based on phylogenomic analysis of universal marker marker genes, offering stronger and orthogonal taxonomic predictions. We found that clade N330390 had significant matches (88.7% average protein identity) with three SPIRE MAGs that were classified as a new, undetermined taxonomic class within the unknown *FCPU426* bacterial phylum. This class was indeed placed close to *Goldbacteria* in the global GTDB phylogeny (Fig 6a), supporting CR-based phylogenetic placement. Conversely, CRs in clade N231481 remained highly divergent even when searched against SPIRE, with an average best-hit protein identity of 57.1% and a maximum of 74.1%. Consistent with the phylogenetic context observed in our CR-based phylogeny, SPIRE’s best hits were mostly represented by MAGs from unclassified *Desulfobacterota_F* and *RBG-13-61-14* phyla (see Supp. Table 2).

## DISCUSSION

Our study unveils the vast hidden biodiversity of CR genes and low-abundance chemosensitive organisms present in natural environments.

From a technical perspective, we proved the effectiveness of our target sequencing strategy in capturing CR genes from across the entire prokaryotic phylogenetic tree, as well as from a variety of environmental samples. Furthermore, target capture sequencing was proven to be an efficient method in complex host-associated microbial habitats, where host DNA tends to be overabundant and bias sequencing results. An example of this is the highly boosted detection of CR genes in the tomato plant phyllosphere included in Pool-1 (see Supp. Fig. 3), where it is usually difficult to deplete plant DNA in order to explore the microbial communities present in plant leaves. On the down side, we found that the performance of capture experiments on heterogeneous sample pools (i.e. from very different environments, as in Pool-1) was lower than that of homogeneous pools. We also faced limitations in attaining the full length of CR genes, as capture probes could only be designed to match the conserved MCPsignal domain. Although we adopted an approach in which we captured and subsequently sequenced DNA fragments up to 1.5 Kbps—enabling the sequencing of off-target regions from the captured CR genes and revealing a broad repertoire of new putative sensing domain families from rare taxa—, the percentage of CR containing LBD sequences obtained was relatively low (6.5%), leaving room for methodological improvement.

Despite such limitations, our analyses demonstrate the existence of a vast, previously overlooked landscape of chemosensitive organisms and chemoreceptor genes in environmental samples. This is illustrated by the fact that the number of unique CR genes identified from the six rhizosphere samples in Pool-2 alone exceeded the total number of CR genes detected in the original 13,000 metagenomic samples from the GMGC catalog used to design the capture panel. We therefore argue that the low CR identification rate estimated from regular shotgun sequencing approaches (7.9 CRs/Gpb) is due to the inability to detect numerous low-abundance chemosensitive microorganisms rather than a lack of biodiversity. Furthermore, the high sequencing depth and coverage obtained for capture-based CR genes in each sample enabled us to detect microorganisms that were undetectable by traditional methods, including standard phylogenetic profiling approaches. Thus, although the MCPsignal domain region of CR genes could not be considered an accurate phylogenetic or taxonomic marker, multi-locus identical and unambiguous mappings against reference genomes proved successful in detecting known plant pathogens and symbionts that were latently present in the rhizosphere.

Notably, because target capture sequencing tolerates sequence divergence of up to 20-40% nucleotide identity^51,52^, our approach allowed us to further discover thousands of unknown CR genes from putative new species. Due to the limitations of MCPsignal domain sequence as a phylogenetic marker, we were unable to infer precise taxonomic assignments for the species carrying such novel CR genes. However, we used phylogenetic reconstruction and cross-validation analysis against the SPIRE database to identify various highly divergent and monophyletic CR clades that strongly support the existence of new bacterial phyla or classes with chemosensing capacity.

We believe that our study not only presents an unprecedented view of the chemoreception in environmental samples, but also opens new avenues for discovering and characterizing novel sensing capabilities of unknown microorganisms. More broadly, we demonstrate that target capture sequencing is a useful tool for exploring functional biodiversity in the rare microbial biosphere.

## METHODS

### Domain extraction

Chemoreceptor genes were extracted using the methodology described in Sanchis-López *et al.*^44^ from all datasets analyzed in this study. Sequences containing the MCPsignal domain (Pfam: PF00015) were retrieved using HMMER v.3.2.1^45,53^ with an e-value of 0.0001. Ligand Binding Domains were also retrieved using Sanchis-López *et al.*^44^ methodology. In brief, since CR sensing regions typically contain ligand-binding domains (LBDs) near the N-terminus, we identified LBDs using three complementary approaches. Putative LBDs were assigned based on: (i) Pfam annotations of conserved ligand-binding domains, (ii) domain localization near the N-terminus, and (iii) sequence analyses of regions lacking clear domain annotations, labeled as “unknown.”

### Sequence retrieving from GMGC catalog

A total of 149,326 CRs sequences were obtained from the GMGCv1^43^ resource (http://gmgc.embl.de), a novel metagenomic catalog combining 84,029 genomes from ProGenomes2^54^ and 302,655,267 unigenes from 13,174 public metagenomic samples from 14 different environments. Sequences containing the MCPsignal domain (Pfam: PF00015) were retrieved using HMMER v.3.2.1^45,53^ with an e-value of 0.0001. Then, LBD characterization was carried out using the pipeline described in Sanchis-López *et al.*^44^. CR sequences catalog and metadata information are available at https://github.com/compgenomicslab/metaCR.

### Ecological annotation of CR sequences

Ecological annotation of the CRs open reading frames (ORFs) was based on the GMGCv1 annotation of the samples where the ORFs were assembled. As for the origin of the samples, we maintained the 14-environment encompassing human-associated (gut, nose, vagina, oral, skin), animal-associated (cat gut, mouse gut, dog gut, pig gut), and environmental (soil, built-environment, marine, wastewater, and freshwater) microbial ecosystems. The number of CRs per habitat was calculated aggregating the information of the samples within each environment.

### Probe set design

The MCPsignal domains of the 149,326 collected sequences were divided into 80-mers (i.e., k-mers of length 80), hereafter probes, with a tiling approach of 20 nt intervals (i.e., 60 nts of overlap) and clustered using CD-HIT v.4.8.1^55^ (with 80% identity and 75% coverage). From 3,249,805 initial k-mers, we obtained 1,354,437 clusters from which we took one representative k-mer as a probe (i.e., 21.4 probes per MCPsignal domain). This set of probes was mapped (using BLASTn from BLAST+ v.2.9.0^56^; min %identity 80%, min %query coverage 90%) to the initial MCPsignal sequences. Further probes were retrieved (same procedure and parameters as above) from regions without any probes mapping to them. To remove promiscuous and nonspecific 80-mers, we mapped our probes to Progenomes2^54^ and to a custom database of organellar genomes using BLASTn once again. Probes that either hit off-target sequences (promiscuous) or a broad range of targets (nonspecific) were discarded. The final set of probes (1,327,258 80-mers) was synthesized as a Roche NimbleGen EZ Developer DNA probe set (Roche Sequencing Solutions, Madison, WI, USA).

### Sample collection and DNA extraction

#### Bulk soil, rhizosphere, and phyllosphere

samples were collected from two different crops, i.e., tomato (*Solanum lycopersicum* L.; greenhouses of HM. Clause Iberica, Almería, Spain) and potato (*Solanum tuberosum* L.; crops from Udapa S.C., Vitoria-Gasteiz, Spain). Bulk soil was sampled in the fraction adjacent to the plant, but not adhered to the roots, and stored at -80°C. Rhizospheric soil was collected from the root, after hand-shaking elimination of the non-adhering soil fraction, by immersion and washing in a solution of PBS and Trition X-100 (0.1%) for 10 min. Then, rhizospheric soil was sieved and collected by centrifugation at 4000 x *g* for 10 min at 4°C. Supernatant was discarded and the rhizospheric soil pellet was stored at -80°C. For the phyllosphere fraction, leaves were collected from the plants, placed in ziplock bags and stored at 4°C for 24 h prior to DNA extraction. DNA from Bulk and Rhizospheric soil was extracted using “DNeasy PowerSoil” (Qiagen, Valencia, CA, USA), following manufacturer’s instructions. DNA from phyllosphere samples was extracted following the protocol described by Suda *et al.*^57^.

#### Lake water

For the lake sample, a volume of 125 mL of water was collected (Casa de Campo Lake, Madrid, Spain) and filtered through a 0.22 µm filter. Then, the filter was stored at -20°C. For DNA extraction, the filter was cut into pieces using sterilized scissors and processed using commercial “DNeasy PowerSoil” (Qiagen, Valencia, CA, USA), following manufacturer’s instructions.

#### Rumen

The rumen sample was collected from a Holstein cow in her first lactation using a custom-built mechanical device to elevate the cow’s snout. Then, approximately 100 mL of ruminal content was extracted by inserting a tube connected to a mechanical pump (Vacubrand ME 2SI, Wertheim, Germany) via the esophagus. The sample was filtered using four layers of sterile cheesecloth to separate the solid elements, and the liquid portion was immediately frozen using liquid nitrogen vapors. These frozen samples were then conveyed to the lab in containers filled with liquid nitrogen and kept at -80°C until analysis. The practice was performed in accordance with EU Directive 2010/63/EU for animal experiments and approved by the ethics committee (NEIKER-OEBA-2017-004). For DNA extraction, the sample was thawed and homogenized using a blender and microbial DNA was extracted using the commercial kit “DNeasy PowerSoil” (Qiagen, Valencia, CA, USA).

#### Human gut

A human fecal sample was collected from an adult asymptomatic individual in December 2019, and ca. 1-2 g were stored within 5 min at -80**°**C. The sample was processed by grinding it to powder with a sterile mortar and pestle under liquid nitrogen. DNA from an aliquot of ca. 1 g was extracted using the AllPrep DNA/RNA Kit (Qiagen) with the following modification: the powder was added to a 2 mL tube pre-filled with 100 μm Zirconium beads (OPS Diagnostics LLC) and 0.6 mL of RLT buffer complemented with 10 μL beta-mercaptoethanol (Sigma-Aldrich) and disrupted twice using the mixer mill Retsch MM400 at 30 Hz for 3 min with 5 min incubations in-between. Cell debris and beads were pelleted by centrifugation at full speed before transferring the supernatant to a DNA-binding column. The remaining steps followed the manufacturer’s instructions. The sampling was reviewed by the Swiss Ethics Committees on research involving humans and was deemed not to require authorization, as it does not fall within the scope of the Human Research Act.

### Samples pooling

Pool-1: This pool included DNA samples from the human gut, rumen, lake, phyllosphere of potato (phyllpot), phyllosphere of tomato (phylltom), three biological replicates of the rhizosphere of potato (rhizpot1, rhizpot2, rhizpot3), three biological replicates of the soil of potato (soilpot1, soilpot2, soilpot3), and three biological replicates of the soil of tomato (soiltom1, soiltom2, soiltom3).

Pool-2: This pool included DNA from three biological replicates of the potato rhizosphere (rhizpot1, rhizpot2, rhizpot3) and three biological replicates of the tomato rhizosphere (rhiztom1, rhiztom2, rhiztom3). All samples in Pool-2 were different from those in Pool-1.

### DNA Fragmentation

DNA fragmentation in Pool-1 was performed using enzymatic digestion NEBNext dsDNA Fragmentase (M0384), resulting in DNA sizes ∼1 Kb. Fragmentation in Pool-2 was done using a Covaris M220 ultrasonicator shearing protocol for 1.5 Kb.

### Target capture experiments

The target capture methodology described below was inspired by the work of Bethune *et al.*^46^, modified as necessary, and applied to both the Pool-1 and Pool-2 experiments.

#### Quality Control and DNA Purification

Fluorometric DNA quantitation was done using a Qubit dsDNA HS Assay Kit (Thermo Fisher Scientific, Waltham, Massachusetts, USA). DNA fragment size distribution and molarity were profiled using the Agilent 4150 TapeStation System (Agilent Technologies, Santa Clara, CA, USA). DNA clean-up and size selection was performed using Agencourt AMPure XP beads (Beckman Coulter).

#### Library Preparation

DNA libraries were prepared according to the NEBNext Ultra II DNA Library Prep Kit for Illumina (E7645, E7103) manual with in-house modifications to obtain longer DNA sizes. The starting amount of fragmented DNA used per sample was 300 ng. All the steps of the manufacturer’s protocol were performed at half the recommended volume to reduce costs^58^. The size selection step was modified to retain the longer sequences, with a single step adding 0.5x AMPure XP beads (Beckman Coulter, Brea, CA, USA) to discard shorter fragments. Samples were indexed using NEBNext Multiplex Oligos for Illumina (Dual Index Primers Set 1, E7600). Thermal cycling parameters for enrichment of Adaptor-ligated DNA were 6 cycles of 130s extension each. Post-PCR clean-up was performed using AMPure XP beads (0.65x). Finally, all samples of each pool were mixed at equimolar concentration for both Pool-1 and Pool-2.

#### Hybridization and enrichment

From the equimolar pools, 1 µg of DNA was taken for capture experiments. Target enrichment experiments were performed according to the Roche NimbleGen Inc. protocol “SeqCap EZ HyperCap Workflow User’s Guide v2.3” with in-house modifications. DNA library pools from the previous step were hybridized with the SeqCap EZ probe set for 24 h at 49.5°C. After washing and recovery of the captured multiplexed DNA sample, amplification of the DNA was done using the Kapa HiFi HotStart ReadyMix with 22 cycles and an extension time of 3 min.

#### Long-read sequencing of Pool-1

The captured and enriched DNA was sequenced in a GridION using a R.9.4.1 flow cell (Oxford Nanopore Technologies, ONT, Oxford, UK). The ligation sequencing kit used was SQK-LSK110, with end prep and adapter ligation performed according to the manufacturer’s instructions. Base-calling was performed using the ONT software Guppy v.4.2.2 in high accuracy mode with quality score (QS) filter > 9 and read length > 50 bp. A total of 5.56 Gbp passed (7,019,121 reads), out of 6.91 Gbp sequenced. N50 of 875 bp.

#### Long-read sequencing of Pool-2

The captured and enriched DNA was sequenced in a MinION using an ONT R.10.4.1 flow cell. The ligation sequencing kit used was SQK-LSK114, with end prep and adapter ligation performed according to the manufacturer’s instructions on the multiplexed NEBNext Ultra II genomic libraries (with Illumina adapters). High accuracy base-calling was performed using the ONT software Guppy v.4.2.2 with filter QS > 15 and read length > 200 bp. A total of 14.93 Gbp passed (10,738,522 reads) out of 17.28 Gbp sequenced. N50 of 1.65 kb.

### Bulk shotgun metagenomics sequencing

DNA samples of Pool-1 and Pool-2 (except human gut) were sequenced with Illumina using Nextera DNA XT libraries in a NovaSeq 6000 platform, PE150 (2 x 150 bp) 8Gbp per sample. Human gut sample (PRJEB90289) was sequenced by Novogene on a NovqSeq 6000 platform, PE150 (2 x 150 bp) at 11Gpb/sample.

### Sequencing data availability

All raw sequencing reads generated in this study will be deposited in the European Nucleotide Archive (ENA) under project accession number PRJEB75710.

### Read processing: trimming, demultiplexing, assembly, and deduplication

#### Assembly of long-read sequences for Pool-1 and Pool-2

Nanopore base-called reads were trimmed with porechop v.0.2.4. Demultiplexing was done with custom scripts, searching for the NEBNext Multiplex Oligos unique sequences (which include Illumina adaptors). A round of BlASTn from BLAST+ v.2.9.0^56^ with the whole index sequence was performed, then the 8 unique nts for each index were identified by text similarity, giving a score to each index per sequence and finally classified based on the best scores and index combination of the samples (pipeline available at https://github.com/compgenomicslab/demultiplex-ont-illumina). In Pool-1, 3,352,198 reads could be demultiplexed, out of the 6,938,358 trimmed. For Pool-2, 9,053,747 reads were correctly demultiplexed, out of 10,738,705 trimmed reads. To remove duplicate reads, representative clusters at 99% identity were identified using CD-HIT v.4.8.1^55^ ( -c 0.99 -aL 0.95 -aS 1).

#### Bulk shotgun metagenomics sequences

Obtained Illumina reads were inspected with FastQC v.0.11.9^59^ and MultiQC v.1.10.1^60^. Reads were trimmed using Trimmomatic v.0.38^61^ (-phred33 ILLUMINACLIP:adapters/NexteraPE-PE.fa:2:30:10 LEADING:3 TRAILING:3 SLIDINGWINDOW:4:15 MINLEN:50). The trimmed Illumina reads were subjected to *de novo* assembly using MEGAHIT v.1.2.8^61,62^ (--presets meta-large). To remove duplicate contigs and to reduce redundancy in the dataset, the assembly output was processed using BBMap v.38.87^63^ dedup.sh (minidentity=99).

#### Host-plant contamination control

For measuring the plant host contamination in the samples sequenced with shotgun metagenomics, the reads were aligned to the reference genomes of tomato (*Solanum lycopersicum* L, GCF_000188115.4) and potato (*Solanum tuberosum* L, GCF_000226075.1) using minimap2 v.2.24-r1122^64^ and SAMtools v.1.12^65^.

### Detection of on-target reads following capture experiments

In order to mine the potential on-target reads/contigs (capture/shotgun metagenomics experiments), first we performed a search against a few MCPsignal representative domains extracted from PfamA-32-A (Supp. materials), using eggNOG-mapper v.5^66^ (-m diamond –itype metagenome). The whole read/contig containing ORFs predicted as potentially on-target were translated into the six possible frames with transeq v.6.5.7^67^ (-frame 6 -clean). Domains were extracted from protein sequences using HMMER v.3.2.1^45,53^ (hmmsearch against Pfam-32-A database). Then, MCPsignal domains were extracted and counted from protein reads, and only one reading frame was kept for each read according to the best MCPsignal hit. In addition, for each on-target read/contig, we extracted Pfam domains and constructed a domain architecture profile based on the two domains immediately before and after the MCPsignal, if present.

To identify unique MCPsignal domains given the ONT reads error rates in Pool-1 and Pool-2, the domains were clustered at 90% and 97% nts identity respectively using cd-hit-est from CD-HIT v.4.8.1^55^ (-aL 0.5 -aS 0.95).

Ligand-binding domains (LBDs) were detected using a combination of Pfam annotations, domain positioning near the N-terminus, and sequence similarity of unannotated N-terminal regions, following the approach outlined in Sanchis-López *et al.*^44^ (See “Domain extraction” section). Only LBDs Pfam signatures identified within open reading frames (ORFs) from the GTDB r207⁴⁸ proteome (see “Mapping to GTDB” section) were retained for downstream analyses.

### Sequencing methods comparison (ONT vs. Illumina)

The on-target ratio was normalized per base pair (bp) by performing ten random subsampling runs, for the same amount of bp given the depth of sequencing for each experiment (40 Mbp Pool-1, 374 Mbp Pool-2). In the shotgun samples, the subsampling was done with seqtk v.1.3^68^, while for the captured demultiplexed pools, subsampling was done with a custom python script. After the subsampling, the starting raw reads were processed for detecting the on-target reads following the same pipeline described above.

### Mapping to GTDB r207

MCPsignal domains (PF00015) were extracted from the proteome of GTDB r207^48^ proteomes using the same approach described in Sanchis-López *et al.*^44,48^ (see “Domain extraction” section). Ligand-binding domains (LBDs) were also retrieved from GTDB ORFs using the same annotation pipeline. Later on, the MCPsignal domain sequence (in nts) was mapped against the target reads (shotgun or capture analyses) using Blastn from BLAST+ v.2.9.0^56^ (-task blastn -evalue 0.0001 -max_target_seqs 10000). The output hits were filtered by sequence identity (100%, ≥ 98%, and ≥ 80%), and by query coverage (≥ 95%). The assignment of a known CR from GTDB was only made when all the hits were 100% identical to the same and unique CR, in order to avoid ambiguity. In case of having more than one unique CR as significant hits, the assignment of the last common ancestor of the CR was calculated over all significant hits taxonomic annotations.

Protein-level comparisons of MCPsignal domains and ligand-binding domains (LBDs) from the captured CRs were performed using DIAMOND blastp v2.0.3^69^ (diamond blastp -e 0.0001 --max-target-seqs 1 --ultra-sensitive --iterate). Significant hits were filtered using a query coverage ≥ 95% and subject coverage ≥ 50%

### Phylogenetic distribution of the known CRs detected

The GTDBr207 tree of bacteria and archaea was concatenated and annotated using ETE Toolkit v.4^70^ and TreeProfiler^71^, and subsequently collapsed to the order level. The sum of the different known CRs detected by order, for each analysis, was calculated and placed in the tree. iTOL^72^ was used for tree visualization.

### Taxonomic profiling of metagenomic samples

mOTUS v.3.0.1^49^ was used to infer species detections on shotgun metagenomics data using default parameters, which required at least three marker genes to be detected to call the presence of a given species (-g3).

### Phylogenetic delineation of novel CRs

The MCPsignal phylogenetic tree was built using protein sequences of the domain extracted from GTDB r207 (305,906 domains), and the MCPsignal domains from shotgun and target capture experiments. A total of 616,738 protein sequences were aligned using FAMSA v.2.2.2^73^, and the phylogenetic tree was reconstructed using FastTree v.2.1.11^74^. Each CR sequence from our experiments was annotated with the protein percentage of identity to the best hit (filtering by query coverage ≥ 95% and subject coverage ≥ 50%) in GTDB r207, as well as the taxonomic information of that best hit. Visualization of the gene tree and the annotations was performed using ETE v.4^70^ and TreeProfiler^71^.

The identification of monophyletic nodes of captured CRs sequences was done using ETE v.4^70^. For each candidate clade, the last common ancestor (LCA) was assigned based on the taxonomic annotation of the nearest GTDB reference sequences in the tree. To ensure robustness, outlier sequences were excluded, and the LCA was only assigned if at least 90% of the reference sequences within the clade shared the same taxonomic label. Additionally, the clade was required to have bootstrap support ≥ 0.8 to be considered for LCA assignment. For comparison, each group of captured sequences was also independently assigned an LCA based on the best-hit taxonomic annotation from GTDB, using the same identity and coverage thresholds described above.

### Mapping to MAGs

To further contextualize captured CRs, sequences containing the MCPsignal domain (Pfam: PF00015) were retrieved using HMMER v.3.2.1^45,53^ with an e-value of 0.0001 from SPIRE database^50^. MCPsignal sequences from capture experiments were mapped using DIAMOND blastp v2.0.3^69^ (diamond blastp -e 0.0001 --max-target-seqs 1 --ultra-sensitive --iterate). Results were filtered using a query coverage ≥ 95% and subject coverage ≥ 50%.

## Acknowledgements (optional).

This study received support by grant PID2021-127210NB-I00 MCIU/AEI/FEDER, UE, National Programme for Fostering Excellence in Scientific and Technical Research, to JHC, CPC and CSL; by grant FPU19/06635 to CSL; by EoI-TSP3-05 (Severo Ochoa CBGP UPM-INIA/CSIC) and a Ramón y Cajal grant (Ref.: RYC2021-034942-I) funded by MCIN/AEI/10.13039/501100011033 and by the European Union “NextGenerationEU”/PRTR to LP; by grant PID2021-125673OB-I00 funded by MICIU/AEI/10.13039/501100011033/ and FEDER “Una manera de hacer Europa”, to ELS; by Severo Ochoa Center of Excellence program SEV-2016-0672, funded by MICIU/AEI/10.13039/501100011033) to JHC and SSH; and by the Personalized Health and Related Technologies (PHRT) grant of the ETH domain (PHRT - 521) to SS. Authors want to thank *HM. Clause Ibérica* (Almería, Spain) and *Udapa S.C.* (Vitoria-Gasteiz, Spain) for granting access to their greenhouses and field plots, respectively, to obtain tomato and potato plants used in this study. We also thank Sandra Nebreda for technical support.

## Author contributions

JHC and ELS conceived the project and provided funding. CSL designed and performed all experiments and computational analyses, generated all figures and tables, and led results interpretation with contributions of JHC. SSH participated in sample processing, setting up and design of capture experiments with contributions of ELS. CPC designed probes for the CRs capture panel. LP participated in the design of the multiplexed long-read capture protocol. SS contributed sample material and funding. OGR contributed sample material. CSL and JHC wrote the manuscript with contributions from all authors.

## SUPPLEMENTARY TABLES

**Supplementary Table 1.**
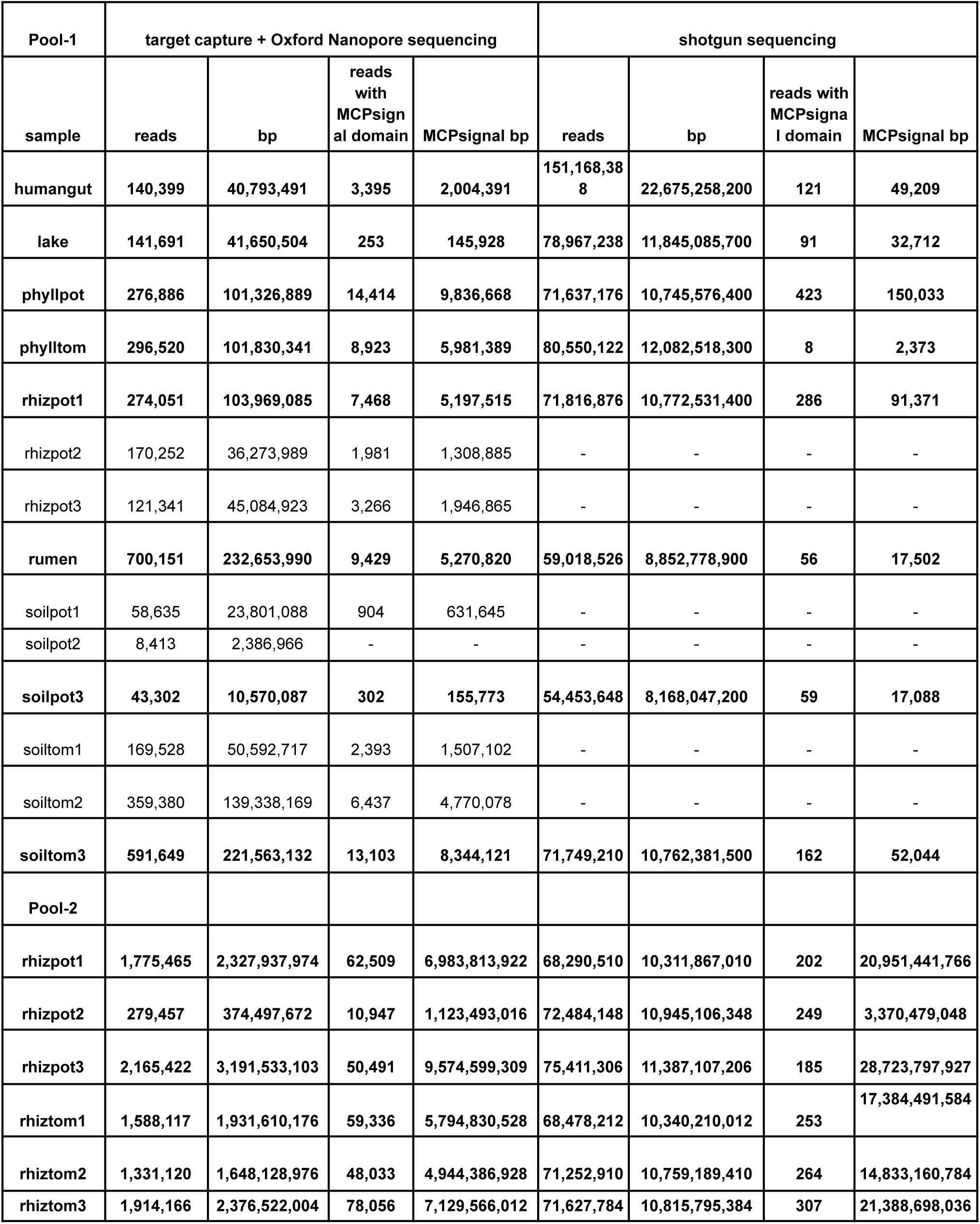
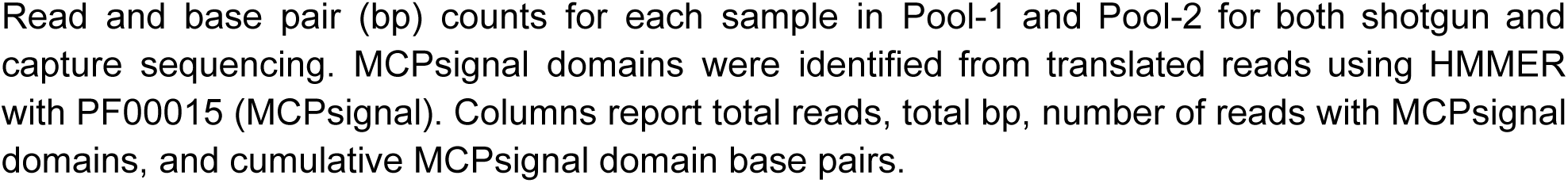
Sequencing and MCPsignal recovery statistics for all capture and shotgun pools.

| Pool-1 | target capture + Oxford Nanopore sequencing |  |  |  | shotgun sequencing |  |  |  |
| --- | --- | --- | --- | --- | --- | --- | --- | --- |
| sample | reads | bp | reads with MCPsignal domain | MCPsignal bp | reads | bp | reads with MCPsignal domain | MCPsignal bp |
| humangut | 140,399 | 40,793,491 | 3,395 | 2,004,391 | 151,168,388 | 22,675,258,200 | 121 | 49,209 |
| lake | 141,691 | 41,650,504 | 253 | 145,928 | 78,967,238 | 11,845,085,700 | 91 | 32,712 |
| phyllpot | 276,886 | 101,326,889 | 14,414 | 9,836,668 | 71,637,176 | 10,745,576,400 | 423 | 150,033 |
| phylltom | 296,520 | 101,830,341 | 8,923 | 5,981,389 | 80,550,122 | 12,082,518,300 | 8 | 2,373 |
| rhizpot1 | 274,051 | 103,969,085 | 7,468 | 5,197,515 | 71,816,876 | 10,772,531,400 | 286 | 91,371 |
| rhizpot2 | 170,252 | 36,273,989 | 1,981 | 1,308,885 | - | - | - | - |
| rhizpot3 | 121,341 | 45,084,923 | 3,266 | 1,946,865 | - | - | - | - |
| rumen | 700,151 | 232,653,990 | 9,429 | 5,270,820 | 59,018,526 | 8,852,778,900 | 56 | 17,502 |
| soilpot1 | 58,635 | 23,801,088 | 904 | 631,645 | - | - | - | - |
| soilpot2 | 8,413 | 2,386,966 | - | - | - | - | - | - |
| soilpot3 | 43,302 | 10,570,087 | 302 | 155,773 | 54,453,648 | 8,168,047,200 | 59 | 17,088 |
| soiltom1 | 169,528 | 50,592,717 | 2,393 | 1,507,102 | - | - | - | - |
| soiltom2 | 359,380 | 139,338,169 | 6,437 | 4,770,078 | - | - | - | - |
| soiltom3 | 591,649 | 221,563,132 | 13,103 | 8,344,121 | 71,749,210 | 10,762,381,500 | 162 | 52,044 |
| Pool-2 |  |  |  |  |  |  |  |  |
| rhizpot1 | 1,775,465 | 2,327,937,974 | 62,509 | 6,983,813,922 | 68,290,510 | 10,311,867,010 | 202 | 20,951,441,766 |
| rhizpot2 | 279,457 | 374,497,672 | 10,947 | 1,123,493,016 | 72,484,148 | 10,945,106,348 | 249 | 3,370,479,048 |
| rhizpot3 | 2,165,422 | 3,191,533,103 | 50,491 | 9,574,599,309 | 75,411,306 | 11,387,107,206 | 185 | 28,723,797,927 |
| rhiztom1 | 1,588,117 | 1,931,610,176 | 59,336 | 5,794,830,528 | 68,478,212 | 10,340,210,012 | 253 | 17,384,491,584 |
| rhiztom2 | 1,331,120 | 1,648,128,976 | 48,033 | 4,944,386,928 | 71,252,910 | 10,759,189,410 | 264 | 14,833,160,784 |
| rhiztom3 | 1,914,166 | 2,376,522,004 | 78,056 | 7,129,566,012 | 71,627,784 | 10,815,795,384 | 307 | 21,388,698,036 |

Read and base pair (bp) counts for each sample in Pool-1 and Pool-2 for both shotgun and capture sequencing. MCPsignal domains were identified from translated reads using HMMER with PF00015 (MCPsignal). Columns report total reads, total bp, number of reads with MCPsignal domains, and cumulative MCPsignal domain base pairs.

**Supplementary Table 2.**
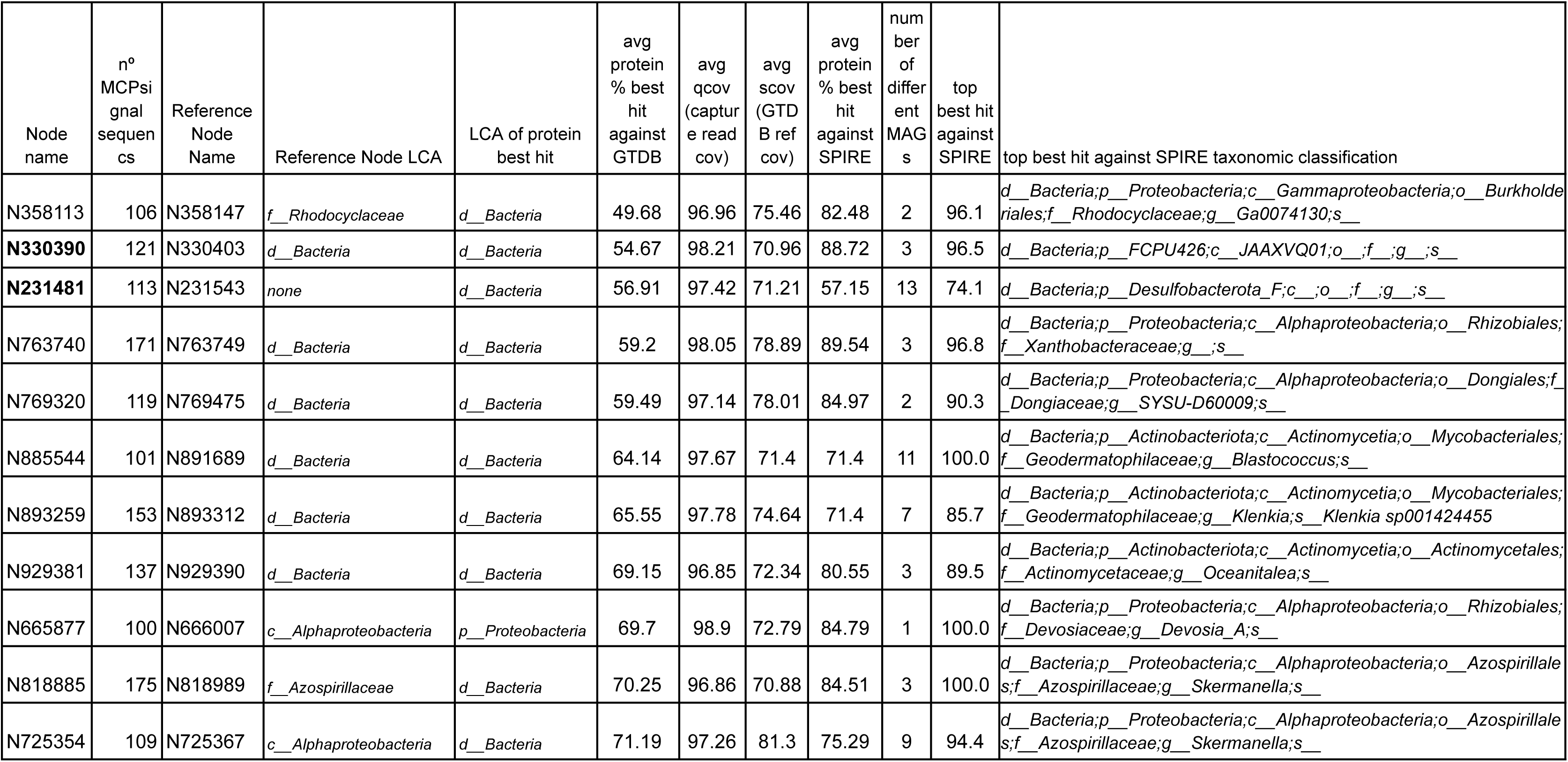
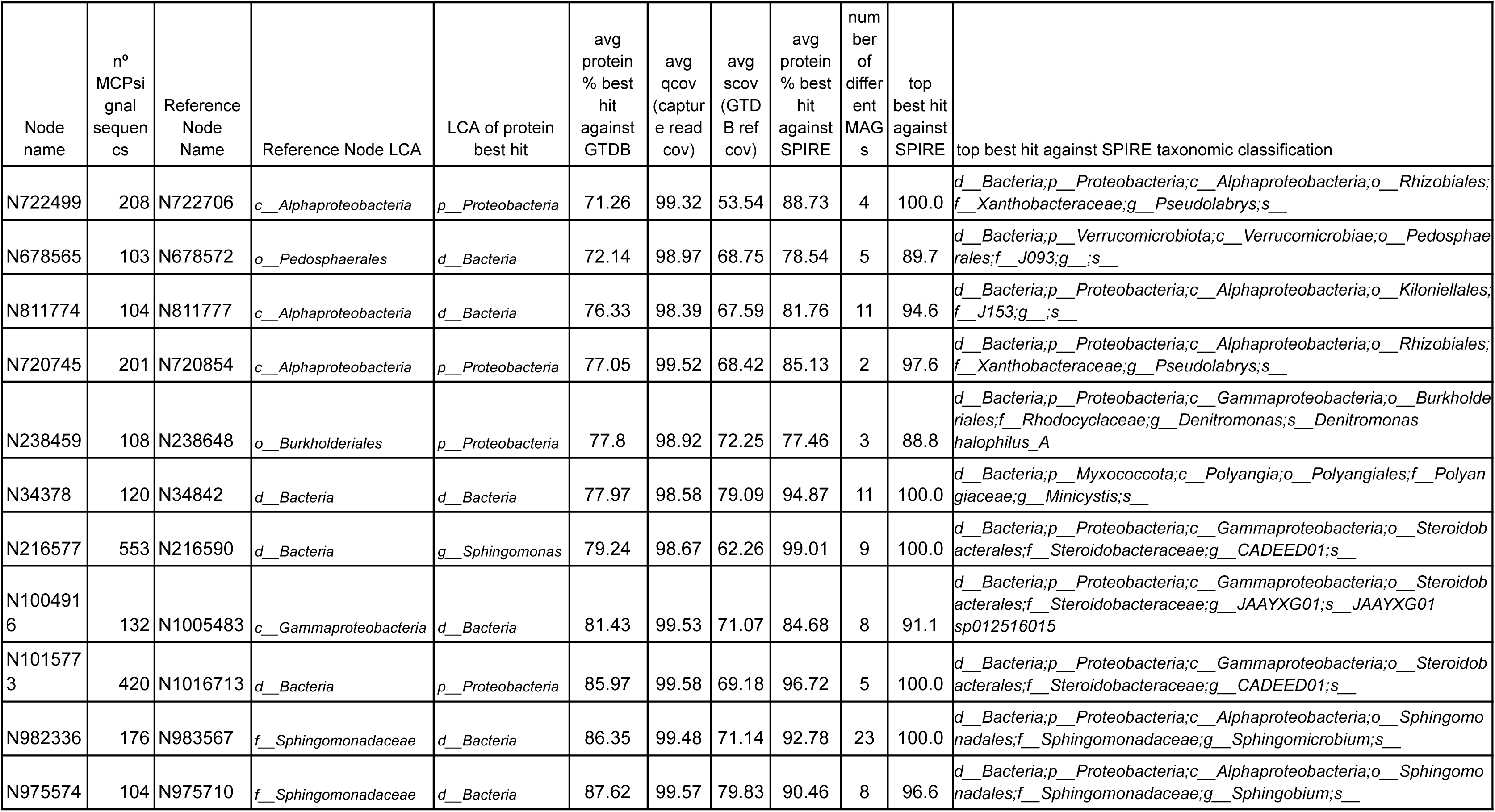
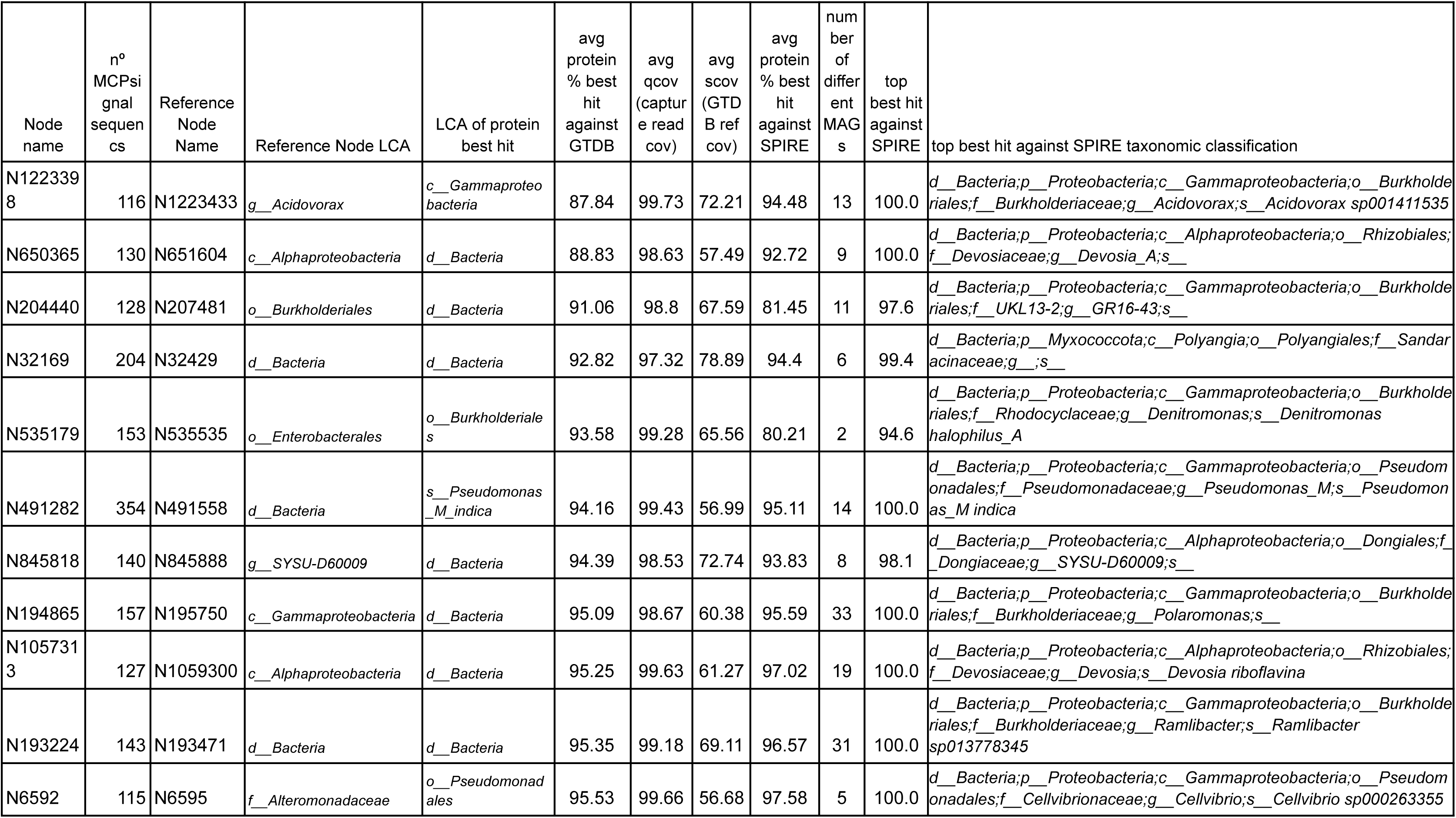
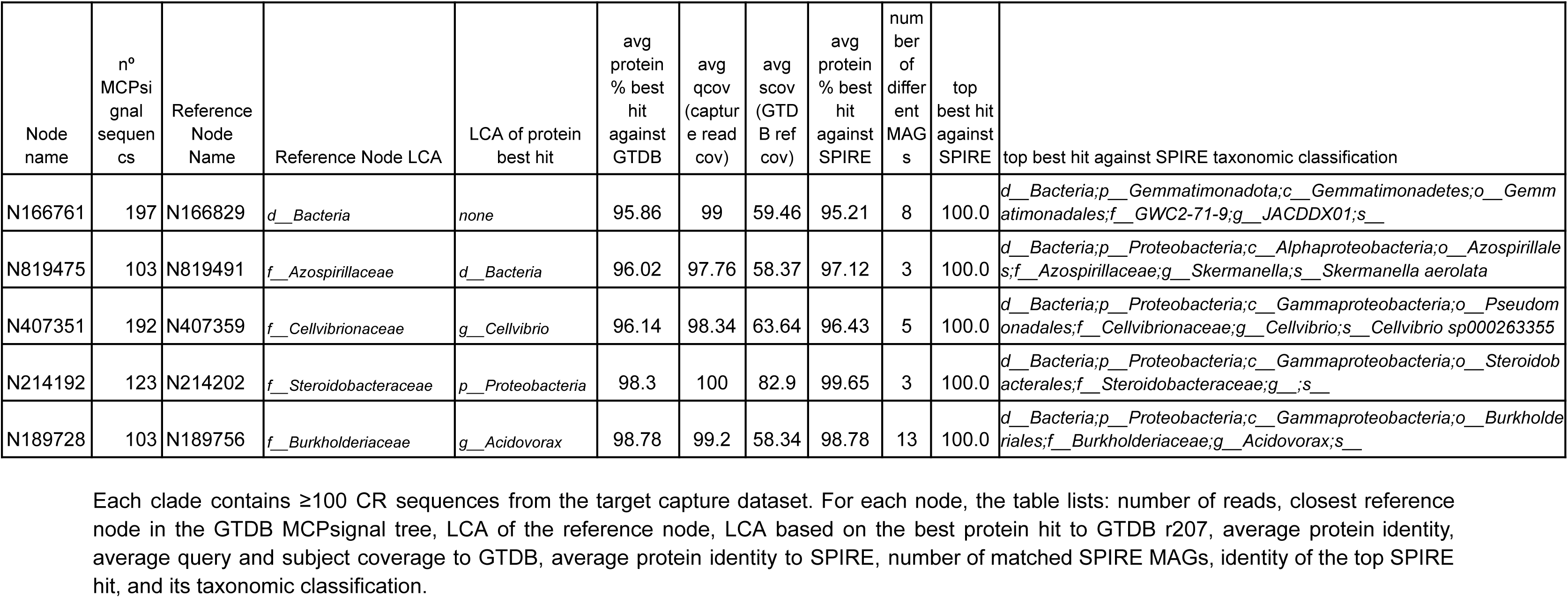
Phylogenetic characterization of novel chemoreceptor clades detected by target capture sequencing.

## SUPPLEMENTARY FIGURES

**Supplementary Figure 1.**
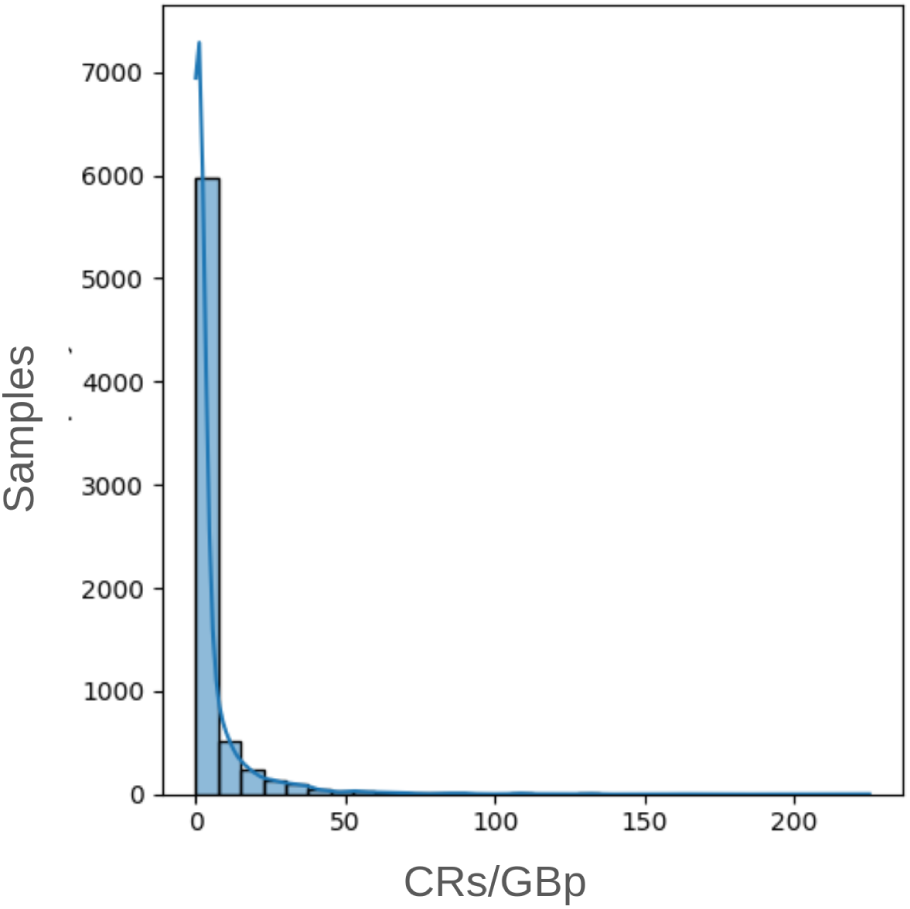
Number of CR genes assembled per Gbp in 13,174 metagenomes from GMGCv1. The histogram represents the distribution of mean CR genes detected per Gbp across the metagenomics samples found in GMGCv1.

**Supplementary Figure 2.**
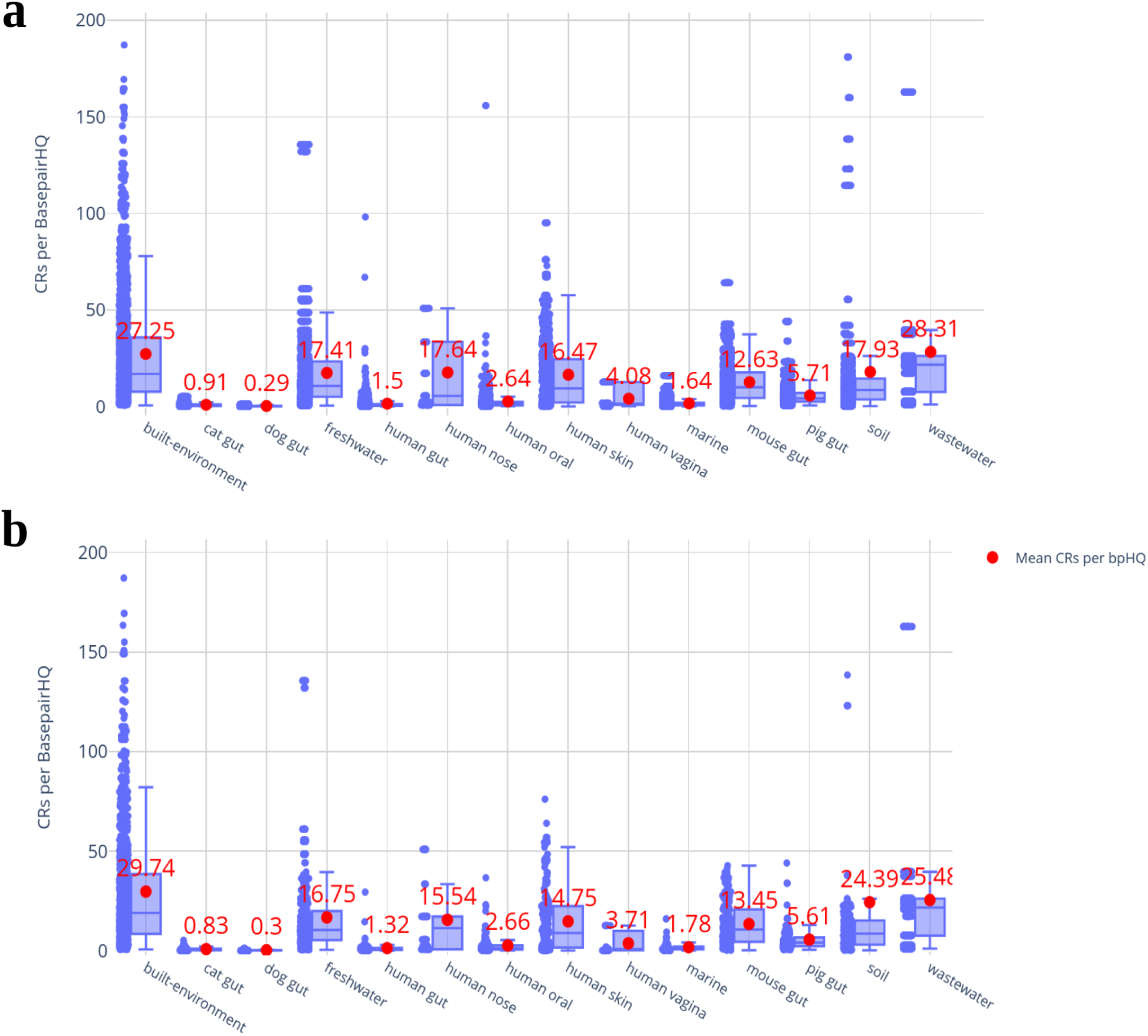
Count normalization of CRs in GMGC catalog. a) CRs per Gbp calculation after subsampling 100 times the same sequencing depth (samples until fulfilling 50 Gbp) in each habitat. b) CRs per Gbp after subsampling 100 times the same number of samples (20) in each habitat. Outliers (>200 CRs/Gbp) are not shown.

**Supplementary Figure 3.**
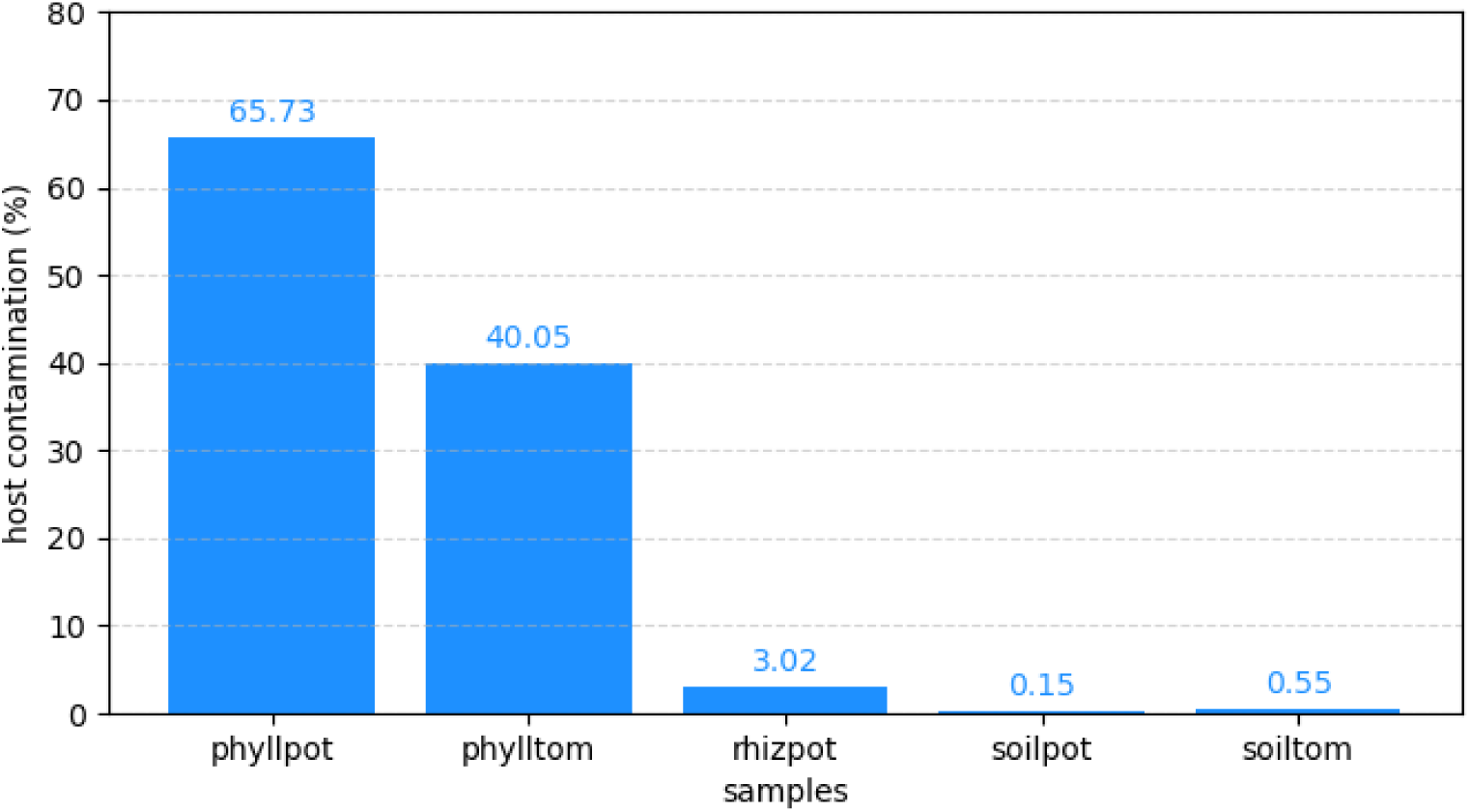
Host-plant contamination rate. The bars show the percentage of reads aligning with the reference genomes of the host plants (tomato, *Solanum lycopersicum* L; potato*, Solanum tuberosum* L) for the samples from Pool-1.

## REFERENCES

1. Keegstra, J. M., Carrara, F. & Stocker, R. The ecological roles of bacterial chemotaxis. Nat. Rev. Microbiol. 20, 491–504 (2022).

2. Raina, J.-B., Lambert, B. S., Parks, D. H., Rinke, C., Siboni, N., Bramucci, A., Ostrowski, M., Signal, B., Lutz, A., Mendis, H., Rubino, F., Fernandez, V. I., Stocker, R., Hugenholtz, P., Tyson, G. W. & Seymour, J. R. Chemotaxis shapes the microscale organization of the ocean’s microbiome. Nature 605, 132–138 (2022).

3. Wuichet, K. & Zhulin, I. B. Origins and diversification of a complex signal transduction system in prokaryotes. Sci. Signal. 3, ra50 (2010).

4. Accessing nutrients as the primary benefit arising from chemotaxis. Curr. Opin. Microbiol. 75, 102358 (2023).

5. Cremer, J., Honda, T., Tang, Y., Wong-Ng, J., Vergassola, M. & Hwa, T. Chemotaxis as a navigation strategy to boost range expansion. Nature 575, 658–663 (2019).

6. Narla, A. V., Cremer, J. & Hwa, T. A traveling-wave solution for bacterial chemotaxis with growth. Proceedings of the National Academy of Sciences 118, e2105138118 (2021).

7. Laganenka, L., Lee, J.-W., Malfertheiner, L., Dieterich, C. L., Fuchs, L., Piel, J., von Mering, C., Sourjik, V. & Hardt, W.-D. Chemotaxis and autoinducer-2 signalling mediate colonization and contribute to co-existence of Escherichia coli strains in the murine gut. Nature Microbiology 8, 204–217 (2023).

8. Hegde, M., Englert, D. L., Schrock, S., Cohn, W. B., Vogt, C., Wood, T. K., Manson, M. D. & Jayaraman, A. Chemotaxis to the Quorum-Sensing Signal AI-2 Requires the Tsr Chemoreceptor and the Periplasmic LsrB AI-2-Binding Protein. J. Bacteriol. (2010). doi:10.1128/jb.01196-10

9. Brumley, D. R., Carrara, F., Hein, A. M., Yawata, Y., Levin, S. A. & Stocker, R. Bacteria push the limits of chemotactic precision to navigate dynamic chemical gradients. Proc. Natl. Acad. Sci. U. S. A. 116, 10792–10797 (2019).

10. Whitchurch, C. B., Leech, A. J., Young, M. D., Kennedy, D., Sargent, J. L., Bertrand, J. J., Semmler, A. B. T., Mellick, A. S., Martin, P. R., Alm, R. A., Hobbs, M., Beatson, S. A., Huang, B., Nguyen, L., Commolli, J. C., Engel, J. N., Darzins, A. & Mattick, J. S. Characterization of a complex chemosensory signal transduction system which controls twitching motility in Pseudomonas aeruginosa. Mol. Microbiol. 52, 873–893 (2004).

11. Colin, R., Ni, B., Laganenka, L. & Sourjik, V. Multiple functions of flagellar motility and chemotaxis in bacterial physiology. FEMS Microbiol. Rev. 45, (2021).

12. Hickman, J. W., Tifrea, D. F. & Harwood, C. S. A chemosensory system that regulates biofilm formation through modulation of cyclic diguanylate levels. Proc. Natl. Acad. Sci. U. S. A. 102, 14422–14427 (2005).

13. Long, Z., Quaife, B., Salman, H. & Oltvai, Z. N. Cell-cell communication enhances bacterial chemotaxis toward external attractants. Sci. Rep. 7, 12855 (2017).

14. Matilla, M. A. & Krell, T. The effect of bacterial chemotaxis on host infection and pathogenicity. FEMS Microbiol. Rev. 42, (2018).

15. Raina, J.-B., Fernandez, V., Lambert, B., Stocker, R. & Seymour, J. R. The role of microbial motility and chemotaxis in symbiosis. Nat. Rev. Microbiol. 17, 284–294 (2019).

16. Stocker, R. & Seymour, J. R. Ecology and physics of bacterial chemotaxis in the ocean. Microbiol. Mol. Biol. Rev. 76, (2012).

17. Buchan, A., LeCleir, G. R., Gulvik, C. A. & González, J. M. Master recyclers: features and functions of bacteria associated with phytoplankton blooms. Nat. Rev. Microbiol. 12, 686–698 (2014).

18. Noman, M., Ahmed, T., Ijaz, U., Shahid, M., Azizullah, Li, D., Manzoor, I. & Song, F. Plant-Microbiome Crosstalk: Dawning from Composition and Assembly of Microbial Community to Improvement of Disease Resilience in Plants. Int. J. Mol. Sci. 22, (2021).

19. Hida, A., Oku, S., Miura, M., Matsuda, H., Tajima, T. & Kato, J. Characterization of methyl-accepting chemotaxis proteins (MCPs) for amino acids in plant-growth-promoting rhizobacterium Pseudomonas protegens CHA0 and enhancement of amino acid chemotaxis by MCP genes overexpression. Biosci. Biotechnol. Biochem. 84, 1948–1957 (2020).

20. Pantigoso, H. A., Newberger, D. & Vivanco, J. M. The rhizosphere microbiome: Plant-microbial interactions for resource acquisition. J. Appl. Microbiol. 133, 2864–2876 (2022).

21. Cerna-Vargas, J. P., Santamaría-Hernando, S., Matilla, M. A., Rodríguez-Herva, J. J., Daddaoua, A., Rodríguez-Palenzuela, P., Krell, T. & López-Solanilla, E. Chemoperception of Specific Amino Acids Controls Phytopathogenicity in Pseudomonas syringae pv. tomato. MBio (2019). doi:10.1128/mbio.01868-19

22. Antúnez-Lamas, M., Cabrera-Ordóñez, E., López-Solanilla, E., Raposo, R., Trelles-Salazar, O., Rodríguez-Moreno, A. & Rodríguez-Palenzuela, P. Role of motility and chemotaxis in the pathogenesis of Dickeya dadantii 3937 (ex Erwinia chrysanthemi 3937). Microbiology 155, 434–442 (2009).

23. Manoharan, L., Kushwaha, S. K., Hedlund, K. & Ahrén, D. Captured metagenomics: large-scale targeting of genes based on ‘sequence capture’ reveals functional diversity in soils. DNA Res. 22, 451–460 (2015).

24. Kolenbrander, P. E., Palmer, R. J., Periasamy, S. & Jakubovics, N. S. Oral multispecies biofilm development and the key role of cell–cell distance. Nat. Rev. Microbiol. 8, 471–480 (2010).

25. Marcos, C. N., Bach, A., Gutiérrez-Rivas, M. & González-Recio, O. The oral microbiome as a proxy for feed intake in dairy cattle. J. Dairy Sci. 107, 5881–5896 (2024).

26. Perry, E. K. & Tan, M.-W. Bacterial biofilms in the human body: prevalence and impacts on health and disease. Front. Cell. Infect. Microbiol. 13, 1237164 (2023).

27. Freter, R., Allweiss, B., O’Brien, P. C., Halstead, S. A. & Macsai, M. S. Role of chemotaxis in the association of motile bacteria with intestinal mucosa: in vitro studies. Infect. Immun. 34, (1981).

28. Butler, S. M. & Camilli, A. Going against the grain: chemotaxis and infection in Vibrio cholerae. Nat. Rev. Microbiol. 3, 611–620 (2005).

29. Pittman, M. S., Goodwin, M. & Kelly, D. J. Chemotaxis in the human gastric pathogen Helicobacter pylori: different roles for CheW and the three CheV paralogues, and evidence for CheV2 phosphorylation. Microbiology 147, 2493–2504 (2001).

30. Alexander, R. P. & Zhulin, I. B. Evolutionary genomics reveals conserved structural determinants of signaling and adaptation in microbial chemoreceptors. Proc. Natl. Acad. Sci. U. S. A. 104, (2007).

31. Ortega, Á., Zhulin, I. B. & Krell, T. Sensory Repertoire of Bacterial Chemoreceptors. Microbiol. Mol. Biol. Rev. 81, (2017).

32. Prevalence and Specificity of Chemoreceptor Profiles in Plant-Associated Bacteria. doi:10.1128/msystems.00951-21

33. Stewart, V. The HAMP signal-conversion domain: static two-state or dynamic three-state? Mol. Microbiol. 91, 853–857 (2014).

34. Ames, P., Zhou, Q. & Parkinson, J. S. HAMP domain structural determinants for signalling and sensory adaptation in Tsr, the Escherichia coli serine chemoreceptor. Mol. Microbiol. 91, 875–886 (2014).

35. Alex Appleman, J. & Stewart, V. Mutational Analysis of a Conserved Signal-Transducing Element: the HAMP Linker of the Escherichia coli Nitrate Sensor NarX. J. Bacteriol. (2003). doi:10.1128/jb.185.1.89-97.2003

36. The superfamily of chemotaxis transducers: From physiology to genomics and back. 45, 157–198 (2001).

37. Sachdeva, R., Campbell, B. J. & Heidelberg, J. F. Rare microbes from diverse Earth biomes dominate community activity. bioRxiv 636373 (2019). doi:10.1101/636373

38. Yooseph, S., Nealson, K. H., Rusch, D. B., McCrow, J. P., Dupont, C. L., Kim, M., Johnson, J., Montgomery, R., Ferriera, S., Beeson, K., Williamson, S. J., Tovchigrechko, A., Allen, A. E., Zeigler, L. A., Sutton, G., Eisenstadt, E., Rogers, Y.-H., Friedman, R., Frazier, M. & Venter, J. C. Genomic and functional adaptation in surface ocean planktonic prokaryotes. Nature 468, 60–66 (2010).

39. Sourjik, V. & Armitage, J. P. Spatial organization in bacterial chemotaxis. The EMBO journal 29, (2010).

40. Jousset, A., Bienhold, C., Chatzinotas, A., Gallien, L., Gobet, A., Kurm, V., Küsel, K., Rillig, M. C., Rivett, D. W., Salles, J. F., van der Heijden, M. G. A., Youssef, N. H., Zhang, X., Wei, Z. & Hol, W. H. G. Where less may be more: how the rare biosphere pulls ecosystems strings. ISME J. 11, 853–862 (2017).

41. Witek, K., Jupe, F., Witek, A. I., Baker, D., Clark, M. D. & Jones, J. D. G. Accelerated cloning of a potato late blight–resistance gene using RenSeq and SMRT sequencing. Nat. Biotechnol. 34, 656–660 (2016).

42. Mamanova, L., Coffey, A. J., Scott, C. E., Kozarewa, I., Turner, E. H., Kumar, A., Howard, E., Shendure, J. & Turner, D. J. Target-enrichment strategies for next-generation sequencing. Nat. Methods 7, 111–118 (2010).

43. Coelho, L. P., Alves, R., Del Río, Á. R., Myers, P. N., Cantalapiedra, C. P., Giner-Lamia, J., Schmidt, T. S., Mende, D. R., Orakov, A., Letunic, I., Hildebrand, F., Van Rossum, T., Forslund, S. K., Khedkar, S., Maistrenko, O. M., Pan, S., Jia, L., Ferretti, P., Sunagawa, S., Zhao, X.-M., Nielsen, H. B., Huerta-Cepas, J. & Bork, P. Towards the biogeography of prokaryotic genes. Nature 601, 252–256 (2022).

44. Sanchis-López, C., Cerna-Vargas, J. P., Santamaría-Hernando, S., Ramos, C., Krell, T., Rodríguez-Palenzuela, P., López-Solanilla, E., Huerta-Cepas, J. & Rodríguez-Herva, J. J. Prevalence and Specificity of Chemoreceptor Profiles in Plant-Associated Bacteria. mSystems 6, e0095121 (2021).

45. Finn, R. D., Coggill, P., Eberhardt, R. Y., Eddy, S. R., Mistry, J., Mitchell, A. L., Potter, S. C., Punta, M., Qureshi, M., Sangrador-Vegas, A., Salazar, G. A., Tate, J. & Bateman, A. The Pfam protein families database: towards a more sustainable future. Nucleic Acids Res. 44, D279–D285 (2015).

46. Bethune, K., Mariac, C., Couderc, M., Scarcelli, N., Santoni, S., Ardisson, M., Martin, J.-F., Montúfar, R., Klein, V., Sabot, F., Vigouroux, Y. & Couvreur, T. L. P. Long-fragment targeted capture for long-read sequencing of plastomes. Appl. Plant Sci. 7, e1243 (2019).

47. Karamitros, T. & Magiorkinis, G. A novel method for the multiplexed target enrichment of MinION next generation sequencing libraries using PCR-generated baits. Nucleic Acids Res. 43, e152 (2015).

48. Parks, D. H., Chuvochina, M., Rinke, C., Mussig, A. J., Chaumeil, P.-A. & Hugenholtz, P. GTDB: an ongoing census of bacterial and archaeal diversity through a phylogenetically consistent, rank normalized and complete genome-based taxonomy. Nucleic Acids Res. 50, D785–D794 (2022).

49. Ruscheweyh, H.-J., Milanese, A., Paoli, L., Karcher, N., Clayssen, Q., Keller, M. I., Wirbel, J., Bork, P., Mende, D. R., Zeller, G. & Sunagawa, S. Cultivation-independent genomes greatly expand taxonomic-profiling capabilities of mOTUs across various environments. Microbiome 10, 212 (2022).

50. Schmidt, T. S. B., Fullam, A., Ferretti, P., Orakov, A., Maistrenko, O. M., Ruscheweyh, H.-J., Letunic, I., Duan, Y., Van Rossum, T., Sunagawa, S., Mende, D. R., Finn, R. D., Kuhn, M., Pedro Coelho, L. & Bork, P. SPIRE: a Searchable, Planetary-scale mIcrobiome REsource. Nucleic Acids Res. 52, D777–D783 (2023).

51. Liu, Y., Johnson, M. G., Cox, C. J., Medina, R., Devos, N., Vanderpoorten, A., Hedenäs, L., Bell, N. E., Shevock, J. R., Aguero, B., Quandt, D., Wickett, N. J., Shaw, A. J. & Goffinet, B. Resolution of the ordinal phylogeny of mosses using targeted exons from organellar and nuclear genomes. Nature Communications 10, 1–11 (2019).

52. Johnson, M. G., Pokorny, L., Dodsworth, S., Botigué, L. R., Cowan, R. S., Devault, A., Eiserhardt, W. L., Epitawalage, N., Forest, F., Kim, J. T., Leebens-Mack, J. H., Leitch, I. J., Maurin, O., Soltis, D. E., Soltis, P. S., Wong, G. K.-S., Baker, W. J. & Wickett, N. J. A Universal Probe Set for Targeted Sequencing of 353 Nuclear Genes from Any Flowering Plant Designed Using k-Medoids Clustering. Syst Biol 68, 594–606 (2018).

53. Potter, S. C., Luciani, A., Eddy, S. R., Park, Y., Lopez, R. & Finn, R. D. HMMER web server: 2018 update. Nucleic Acids Res. 46, W200–W204 (2018).

54. Mende, D. R., Letunic, I., Maistrenko, O. M., Schmidt, T. S. B., Milanese, A., Paoli, L., Hernández-Plaza, A., Orakov, A. N., Forslund, S. K., Sunagawa, S., Zeller, G., Huerta-Cepas, J., Coelho, L. P. & Bork, P. proGenomes2: an improved database for accurate and consistent habitat, taxonomic and functional annotations of prokaryotic genomes. Nucleic Acids Res. 48, D621–D625 (2020).

55. Fu, L., Niu, B., Zhu, Z., Wu, S. & Li, W. CD-HIT: accelerated for clustering the next-generation sequencing data. Bioinformatics 28, 3150 (2012).

56. Camacho, C., Coulouris, G., Avagyan, V., Ma, N., Papadopoulos, J., Bealer, K. & Madden, T. L. BLAST+: architecture and applications. BMC Bioinformatics 10, (2009).

57. Suda, W., Oto, M., Amachi, S., Shinoyama, H. & Shishido, M. A Direct Method to Isolate DNA from Phyllosphere Microbial Communities without Disrupting Leaf Tissues. Microbes Environ. 23, 248–252 (2008).

58. Hale, H., Gardner, E. M., Viruel, J., Pokorny, L. & Johnson, M. G. Strategies for reducing per-sample costs in target capture sequencing for phylogenomics and population genomics in plants. Appl. Plant Sci. 8, e11337 (2020).

59. Babraham Bioinformatics - FastQC A Quality Control tool for High Throughput Sequence Data. at <www.bioinformatics.babraham.ac.uk/projects/fastqc/>

60. Ewels, P., Magnusson, M., Lundin, S. & Käller, M. MultiQC: summarize analysis results for multiple tools and samples in a single report. Bioinformatics 32, 3047–3048 (2016).

61. Bolger, A. M., Lohse, M. & Usadel, B. Trimmomatic: a flexible trimmer for Illumina sequence data. Bioinformatics 30, 2114 (2014).

62. Li, D., Liu, C.-M., Luo, R., Sadakane, K. & Lam, T.-W. MEGAHIT: an ultra-fast single-node solution for large and complex metagenomics assembly via succinct de Bruijn graph. Bioinformatics 31, 1674–1676 (2015).

63. Bushnell, B. BBMap: A Fast, Accurate, Splice-Aware Aligner. (Lawrence Berkeley National Lab. (LBNL), Berkeley, CA (United States), 2014). at <https://www.osti.gov/servlets/purl/1241166>

64. Li, H. Minimap2: pairwise alignment for nucleotide sequences. Bioinformatics 34, 3094–3100 (2018).

65. Danecek, P., Bonfield, J. K., Liddle, J., Marshall, J., Ohan, V., Pollard, M. O., Whitwham, A., Keane, T., McCarthy, S. A., Davies, R. M. & Li, H. Twelve years of SAMtools and BCFtools. Gigascience 10, giab008 (2021).

66. Huerta-Cepas, J., Szklarczyk, D., Heller, D., Hernández-Plaza, A., Forslund, S. K., Cook, H., Mende, D. R., Letunic, I., Rattei, T., Jensen, L. J., von Mering, C. & Bork, P. eggNOG 5.0: a hierarchical, functionally and phylogenetically annotated orthology resource based on 5090 organisms and 2502 viruses. Nucleic Acids Res. 47, D309–D314 (2018).

67. Madeira, F., Pearce, M., Tivey, A. R. N., Basutkar, P., Lee, J., Edbali, O., Madhusoodanan, N., Kolesnikov, A. & Lopez, R. Search and sequence analysis tools services from EMBL-EBI in 2022. Nucleic Acids Res. 50, W276–W279 (2022).

68. GitHub - lh3/seqtk: Toolkit for processing sequences in FASTA/Q formats. GitHub at <https://github.com/lh3/seqtk>

69. Buchfink, B., Xie, C. & Huson, D. H. Fast and sensitive protein alignment using DIAMOND. Nature Methods 12, 59–60 (2014).

70. Huerta-Cepas, J., Serra, F. & Bork, P. ETE 3: Reconstruction, Analysis, and Visualization of Phylogenomic Data. Mol. Biol. Evol. 33, 1635–1638 (2016).

71. Deng, Z., Hernández-Plaza, A. & Huerta-Cepas, J. TreeProfiler: A command-line tool for computing and visualizing phylogenetic profiles against large trees. bioRxiv 2023.09.21.558621 (2023). doi:10.1101/2023.09.21.558621

72. Letunic, I. & Bork, P. Interactive Tree of Life (iTOL) v6: recent updates to the phylogenetic tree display and annotation tool. Nucleic Acids Res. gkae268 (2024).

73. Deorowicz, S., Debudaj-Grabysz, A. & Gudyś, A. FAMSA: Fast and accurate multiple sequence alignment of huge protein families. Sci. Rep. 6, 1–13 (2016).

74. Price, M. N., Dehal, P. S. & Arkin, A. P. FastTree 2 – Approximately Maximum-Likelihood Trees for Large Alignments. PLoS One 5, e9490 (2010).

